# Robust cellular transformations of PET deconstruction products by import of glycol esters

**DOI:** 10.1101/2025.10.01.679844

**Authors:** Roman M. Dickey, Esun Selvam, Erha Andini, Priyanka Nain, Dionisios G. Vlachos, Aditya M. Kunjapur

## Abstract

Efforts to transform polyethylene terephthalate (PET) deconstruction products using live cells have been limited by terephthalic acid (TPA) uptake. Here, we used an intracellular carboxylate reduction assay to show that apparent TPA uptake in *E. coli* cells that lack a dedicated TPA transporter sharply increases between pH 5-6. Furthermore, we discovered that glycol ester deconstruction products, mono(2-hydroxyethyl) terephthalate (MHET) and bis(2-hydroxyethyl) terephthalate (BHET), surprisingly each result in rapid pH-independent uptake. We exploited glycol ester uptake along with deletion of 22 cellular oxidoreductases to design intracellular hydrolysis routes for synthesis of upcycled reduction products from BHET at >90% yields, and from real PET wastes after tandem catalytic glycolysis and cell-based valorization at >80% combined yields. Our work has important ramifications for PET utilization by cells and adds new perspectives on the evolution of the PETase/MHETase system.

## Introduction

While society relies on synthetic polymers, the growth in plastic waste is a global environmental problem that is estimated to reach 121 million metric tons by 2050 without intervention^1^. Polyethylene terephthalate (PET) is an exemplary plastic that is one of the most abundantly produced polyester and is widely applied in single use. Chemical recycling of PET via catalytic or enzymatic routes can completely hydrolyze all ester bonds, forming terephthalic acid (TPA) and ethylene glycol (EG), or generate alternative deconstruction products, bis(2-hydroxyethyl) terephthalic acid (BHET) and mono(2-hydroxyethyl) terephthalic acid (MHET) (**Fig 1A**). In the context of PET recycling or upcycling, after the discovery of a PET hydrolase (PETase) and a MHET hydrolase (MHETase) from *Ideonella sakaiensis*, the biocatalysis community has largely focused on discovering or engineering enzymes that catalyze complete deconstruction of PET to TPA and EG with increased efficiency^2^. A smaller number of efforts have reported engineering assimilation or valorization of TPA or EG by microbial cells^3–7^; however, the use of cell-based biocatalysts for deconstruction or transformation of deconstructed PET has lagged considerably behind chemical or enzymatic alternatives^8^. This is despite the ability of whole cells to facilitate a larger number of valorizing transformations while also obviating the need for cell lysis and enzyme purification, thus potentially lowering catalyst costs at industrial scales. Given the polymeric nature of the feedstock, some initial deconstruction steps must occur in an extracellular environment; however, we sought to critically examine the conventional notion that complete extracellular hydrolysis to TPA and EG would be beneficial for cellular transformations.

**Fig. 1.**
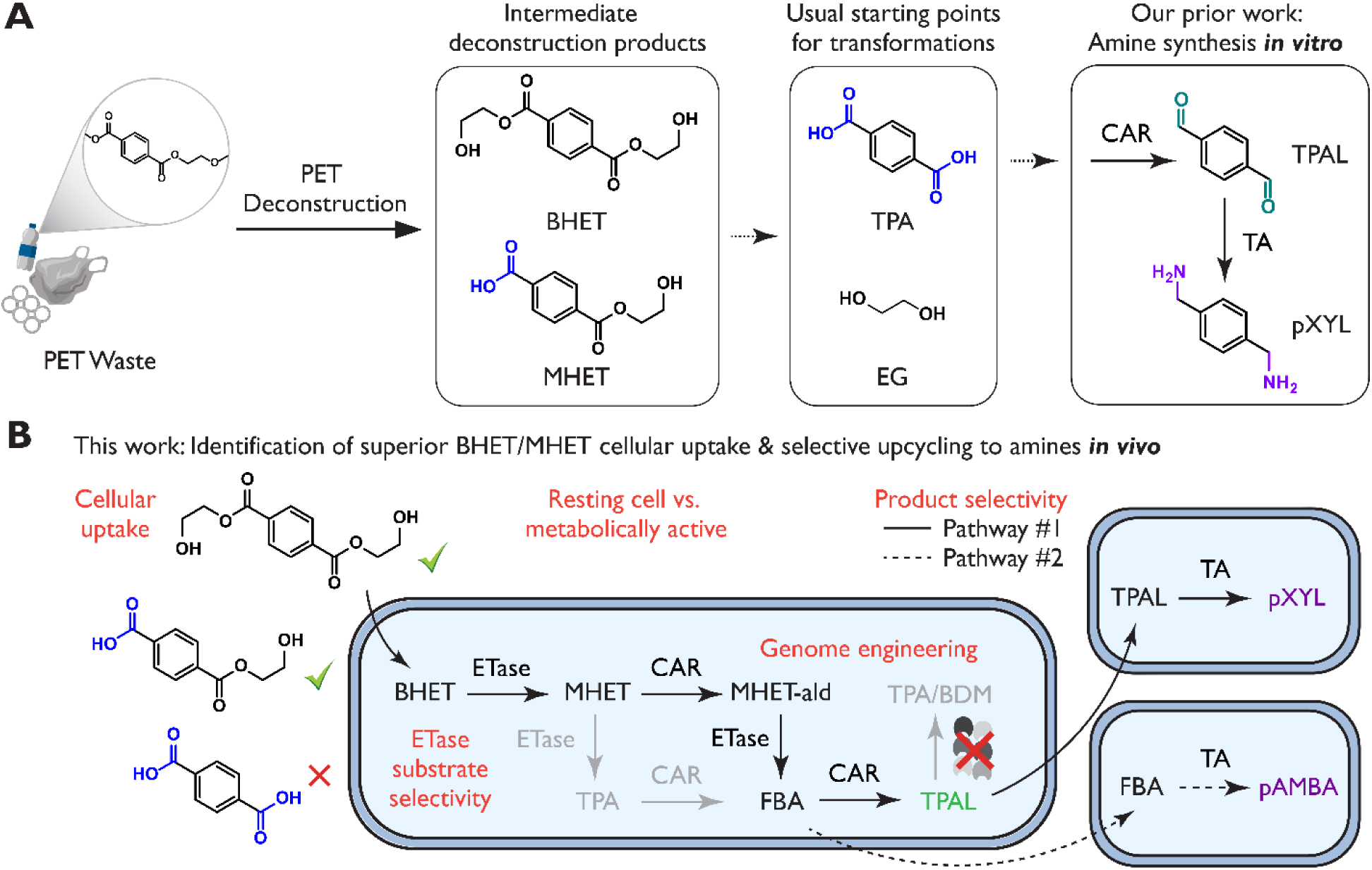
The strategy for biologically upcycling PET using cells based on alternative deconstruction products that exhibit better uptake. **(A)** The complete and alternative deconstruction products of PET shown in the context of our prior upcycling work performed *in vitro*. **(B)** Summary of topics and cellular transformation routes investigated in this work, culminating in high-yield biosynthesis of TPAL, pAMBA, or pXYL from real PET wastes after catalytic glycolysis.

A key bottleneck in these processes is slow cellular uptake of TPA. In bioconversion and upcycling strategies, TPA is commonly prioritized due to its carbon richness compared to EG and potential to serve as a precursor for value-added aromatic compounds. However, as TPA cannot readily diffuse through microbial cell membranes at neutral pH, the expression of TPA transporter genes is thought to be essential to TPA catabolism or upcycling^9^. Thus, the design of cell-based TPA transformations typically involves the expression and engineering of an active TPA transporter or less often methods to alter cell permeability. In *Pseudomonas putida*, heterologous expression of *tpaK*, which encodes an MFS-type TPA transporter from *Rhodococcus jostii*, was used to achieve β-ketoadipic acid production from catalytically depolymerized PET^4^. Interestingly, the authors observed that secretion tags were not required for BHET conversion, speculating that either the PETase/MHETase might exit cells, that BHET might enter cells passively, or that a native transporter might allow BHET uptake. Given the prominence of *Escherichia coli* in synthetic biology and industrial processes, it is an attractive host for TPA transformations or utilization. Recently, adaptive laboratory evolution (ALE) of *E. coli* with heterologous expression of the TpaK transporter from *R. jostii* and genes for TPA assimilation reinforced the notion that TPA transport is limiting even after heterologous expression of transporters^10^. To avoid active TPA transport, alternative approaches to enhance cell membrane permeability and/or TPA uptake have been employed including modification of membrane proteins^11^, lowering culture pH (5.5) or using chemical additives^12^. However, these strategies can decrease downstream pathway activity cell viability, limiting their practical relevance^13^.

We wondered if we could avoid the challenge of TPA uptake by instead targeting the alternative deconstruction products, the glycol esters MHET and BHET, as starting substrates for intracellular transformations (**Fig. 1B**). We used a carboxylic acid reductase (CAR) to modify the carboxylates present in both TPA and MHET as a proxy for detecting cellular uptake and to produce terephthalaldehyde (TPAL), which is a valuable monomer with uses in thermosets^14–16^, polyspiroacetals^17^, polymeric organic frameworks^18^, nanocomposites^19^, and other polymer reactions^20–23^, as well as in the production of new Schiff Bases ligands^24–26^. The reactive aldehyde functionality of TPAL also enables potential biosynthetic routes to numerous alternative chemistries^27–29^, including to the valuable diamine *para*-xylylenediamine (PXDA or pXYL), which has an estimated market size of USD ∼1 billion^30–35^.

Here, we report the enhanced uptake of BHET and MHET, and we exploit this finding for the design of whole cell cascades capable of upcycling PET deconstruction products at neutral pH. We show that MHET is efficiently transported intracellularly and converted toward its aldehyde (2-hydroxyethyl 4-formylbenzoate, or MHET-ald) at neutral and acidic pH; however, TPA uptake and conversion toward its corresponding aldehydes (4-formylbenzoic acid, FBA, and TPAL) is strictly limited to acidic conditions (pH < 6.2). To enable intracellular hydrolysis of BHET and MHET for subsequent transformations, we screen published PETases and cutinases for general intracellular “ETase” activity under ambient conditions. We then show that coupling our selected PETase and CAR enzymes within the same cell enables the biosynthesis of TPAL at high yields. Finally, we show that this PETase-CAR system can be coupled with an ω-transaminase for high-yield biosynthesis of pXYL from products of catalytic glycolysis using real PET textile and bottle waste streams. Overall, our work has significant ramifications for those interested in supplying PET deconstruction products to cellular catalysts, whether for valorization or for catabolism.

## Results

### Cellular uptake of TPA is controlled by pH

To measure apparent uptake of carboxylate group containing PET deconstruction products into *E. coli*, we created an enzyme-coupled assay based on heterologous expression of a CAR shown *in vitro* to possess activity on our candidate substrates (CAR from *Mycobacterium avium,* MaCAR)^36^ (**Fig. 2A**). To retain desired aldehyde products for eventual upcycling reactions, we expressed MaCAR in the previously engineered *E. coli* RARE.Δ16 strain designed for TPAL retention^37^. We supplemented TPA, MHET, as well as mono-substituted carboxylates FBA (the single reduction product of TPA) and 4-hydroxymethylbenzoic acid (HMBA, a potential over-reduction product), to cells expressing MaCAR at a starting pH of 7.5 under aerobic culturing conditions. To ensure that all potential intermediates are collected in the supernatant, we performed a methanol-based extraction of all collected samples. We observed full conversion of MHET, FBA, and HMBA to their respective aldehydes after 4 h, indicating these substrates are readily imported by cells **(Fig. 2B)**. However, we observed no conversion of the dicarboxylic acid TPA, consistent with this step being a bottleneck. Interestingly, we did observe the full conversion of TPA after 24 h, suggesting a delayed uptake relative to monocarboxylic acids. Given that others have briefly observed that pH can play a role in TPA uptake^11^, we monitored culture pH alongside TPA conversion as a function of time. We found that pH decreases correlated to TPA conversion **(Fig. 2C)**, suggesting a simple explanation for the delay. To collect further supporting evidence, we saw that buffering the culture media abrogated TPA conversion **(Fig. S1)**, and that lowering initial culture pH accelerated TPA uptake and conversion **(Fig. 2D)**.

**Fig. 2.**
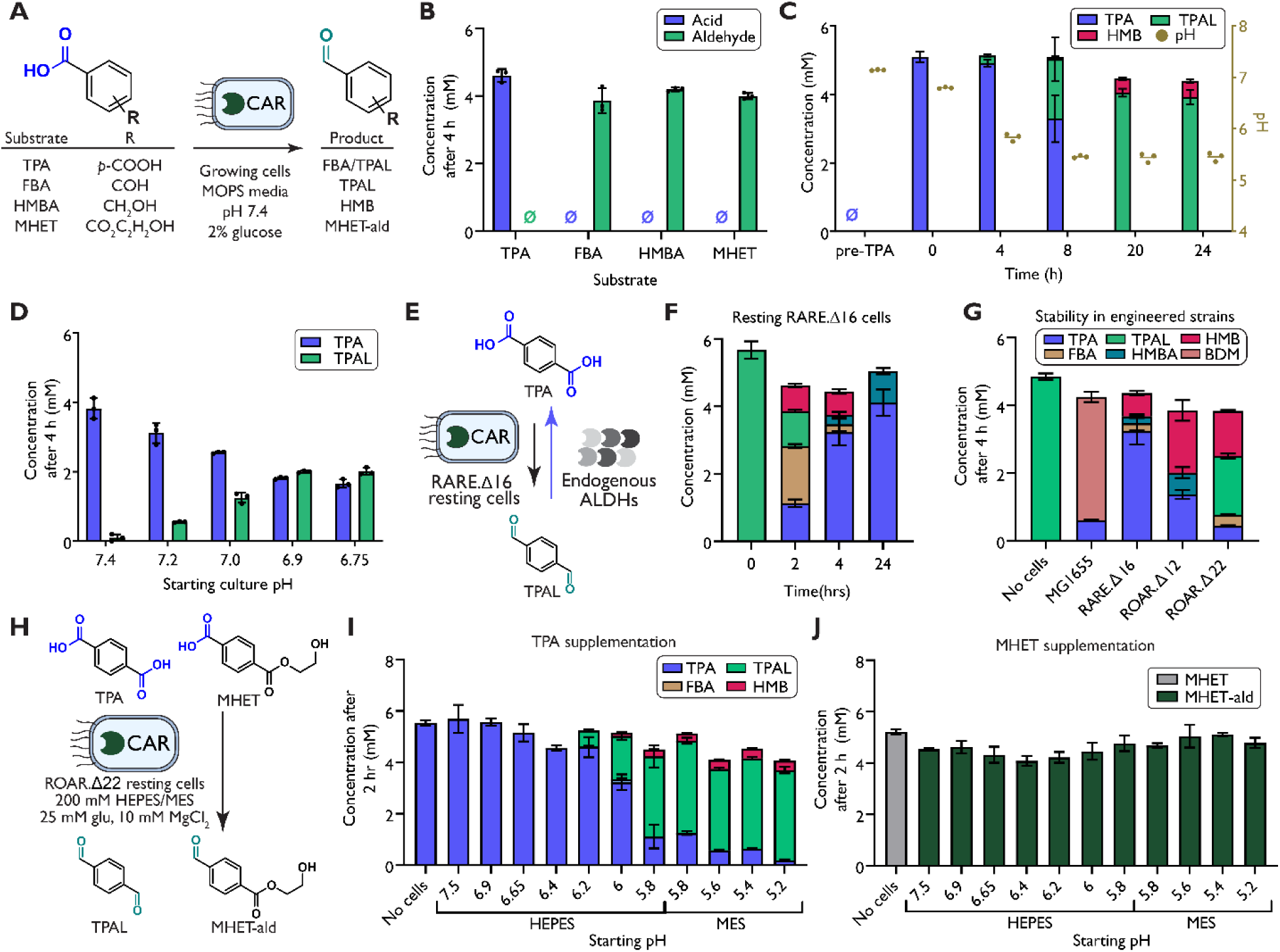
The influence of pH on apparent cellular uptake of TPA and MHET. (**A**) Experimental design, which featured cells of the *E. coli* RARE.Δ16 strain expressing MaCAR cultured in MOPS media with 10 mM MgCl_2_ and 2 % glucose at pH of 7.4. Cells were induced at mid-exponential phase (OD_600_= 0.5-0.7), dropped to 30°C, and supplemented with 5 mM of carboxylic acid substrate. (**B**) Endpoint assay showing the conversion of acid substrates to their corresponding aldehydes after 4 h. (**C**) Time course of the relationship between pH and TPAL production from supplemented TPA. (**D**) Variation of initial pH, which led to conversion of TPA to TPAL at 4 h. Starting pH was measured before the addition of 5 mM TPA. (**E**) Conceptual illustration showing that endogenous aldehyde dehydrogenases (ALDHs) can mask TPA conversion in resting RARE.Δ16 cells. (**F**) Time course of supplemented TPAL oxidation in resting RARE.Δ16 cells. (**G**) The newly engineered *E. coli* ROAR.Δ22 strain limits oxidation and reduction of TPAL compared to wild-type (MG1655) and previously engineered strains (RARE.Δ16 and ROAR.Δ12). (**H**) ROAR.Δ22 cells with MaCAR enable efficient generation and stability of aldehydes (TPAL from TPA and MHET-ald from MHET) to determine uptake of (**I**) TPA and (**J**) MHET at pH ranges from 4.8-7.5. Samples sizes are *n* = 3 using biological replicates. Data shown are mean ± s.d.

To more extensively study how pH affects TPA and MHET import, we transitioned from growing cells to resting cells, allowing for a more controlled reaction. However, we found that resting cells of the engineered RARE.Δ16 host rapidly oxidized TPAL, which is a phenomenon not present in growing cells and could counteract carboxylate reduction (**Fig. 2E-F**). To negate this, we further engineered the RARE.Δ16 to include knockouts of 6 aldehyde dehydrogenases that have been shown to mitigate aryl aldehyde oxidation^38^, generating the ROAR.Δ22 strain. We found that the ROAR.Δ22 host increased TPAL (4.2-fold) and MHET-ald (3.7-fold) concentrations compared to RARE.Δ16 after 4 h (**Fig. 2G**, **Fig. S2**). The strain exhibited negligible fitness difference nor sensitivity of aldehyde retention in various reaction media (pH and glucose concentration) (**Fig. S2-5**). We resuspended resting cells of ROAR.Δ22 that expressed MaCAR with TPA or MHET at starting pH ranging from 4.8-7.5 in increments of 0.2 (**Fig. 2H**). As expected, TPA conversion increased as we lowered pH, until TPA precipitated out of solution at pH 5.0 and 4.8 **(Fig. 2I)**. We could only observe TPA conversion at pH 6.2 or lower, with a maximum conversion of 92% at pH 5.2, indicating that TPA uptake can occur in acidic pHs. The observed TPA uptake correlates well with the predicted formation of the partially deprotonated TPA, highlighting the possibility that the mono-protonated form of TPA is accepted by native transporters (**Fig. S6**). Excitingly, we observed the full conversion of MHET towards MHET-ald at all pH tested **(Fig. 2J)**, and we sought to investigate the design of new transformation routes that would exploit this finding.

### Screening of “ETase” activity to enhance cellular uptake of PET deconstruction products

Because these experiments showed that MHET uptake was fast and unaffected by reaction pH, we hypothesized that intracellular expression of selective esterases (here designated as “ETases” for potential activity on PET, BHET, MHET, and/or MHET-ald) could facilitate complete ester hydrolysis after uptake **(Fig. 3A)**. We desired an ETase that had activity on BHET, MHET and MHET-ald at 30°C for intracellular TPA production with limited membrane leakage that could lead to TPA production extracellularly^39,40^. We screened 10 previously engineered ETases on BHET, MHET and MHET-ald. We selected five engineered variants of the *Ideonella sakaiensis* PETase (FAST^41^, HOT^42^, Dura^43^, Themo-Stable: denoted here as Thermo^44^ and TS^45^) two leaf-branch compost cutinase variants (LCC^39^, LCC-ICCG: denoted here as ICCG^46^), two variants of bacterium HR29 PETase (BhrPETase^47^ and Turbo^48^) and one metagenome-derived PETase (PES-H1:denoted here as PES^49^). While many of these enzymes were primarily studied at elevated temperatures for PET deconstruction, we sought to measure ETase activity under ambient conditions (**Table S1**).

**Fig. 3.**
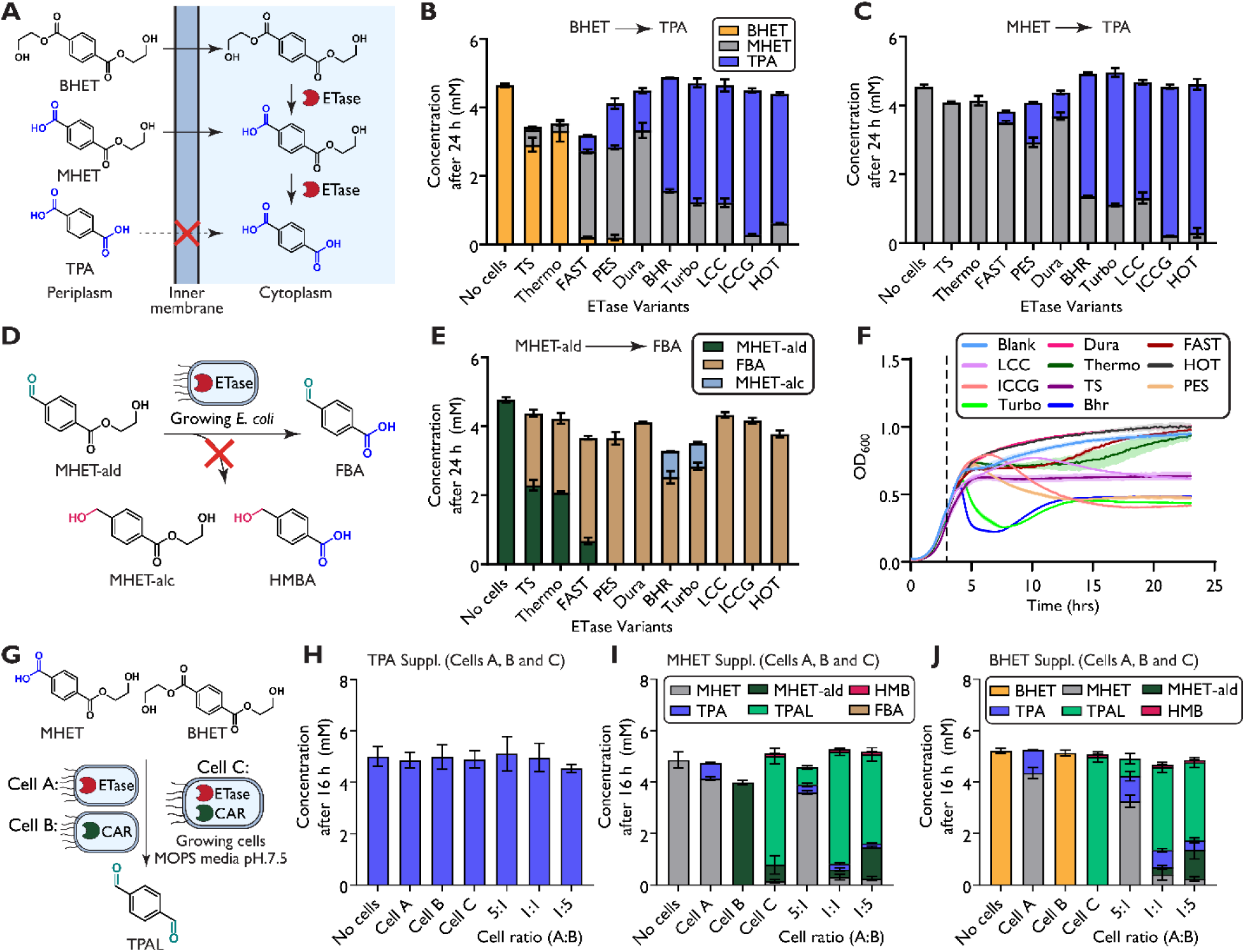
Evaluating ETase specificity on PET deconstruction products for intracellular hydrolysis. (**A**) Intercellular expression of ETases could allow for TPA to be produced within the cell, preventing the need for active transport. Supplementation of 5 mM (**B**) BHET (**C**) MHET or (**E**) MHET-ald to RARE.Δ16 cell expressing ETase variants in MOPS media. Endpoint concentrations were measured after 24 h. (**D**) The conversion of a novel substate, MHET-ald, to FBA. (**F**) Effect of ETase expression (induction indicated by dotted line at 3 h) on cellular growth. (**G**) Evaluation of coupling FAST and MaCAR in growing cells in either separate strains (Cell A and B respectively) or in the same strain (Cell C) for the production of TPAL from (**H**) TPA, (**I**) MHET or (**J**) BHET. Samples sizes are *n* = 3 using biological replicates. Data shown are mean ± s.d.

We expressed each ETase initially in RARE.Δ16 in growing cells to track cell viability and to test activity on supplemented BHET, MHET and MHET-ald. We observed a wide range of TPA production when cultures were supplemented with BHET and MHET, indicating differences in substrate specificity and activity between engineered variants, as well as the robust uptake of BHET **(Fig. 3B-C)**. Analysis of the ETase activity allowed us to characterize several BHET-selective ETases, such as FAST, PES and DURA, which had high BHET conversion and low MHET conversion. We also observed several generalist ETases such as BHR, Turbo, LCC, ICCG, and HOT, with high conversion of both BHET and MHET. Surprisingly, we noticed that our entire library of ETases had activity on MHET-ald (**Fig. 3D-E**), and obtained higher conversions on MHET-ald than MHET **(Fig. S7-8)**. After profiling ETase specificity, we noticed a subset of ETases has substantial effects to overexpression (BHR, Turbo, ICCG and PES) with others having minor effects (Thermo, TS, LCC, Dura, HOT and FAST) **(Fig. 3F)**. Considering both activity and toxicity, FAST showed promise for further applications.

### Coupling ETase and CAR to explore the enhanced uptake of BHET and MHET

After identification of viable ETases, we reasoned that by expressing FAST and MaCAR together in one-pot we could produce TPAL from BHET and MHET at neutral pH in growing cells (**Fig. 3G**). We compared expression of FAST and MaCAR in separate cells (Cell A and Cell B, respectively) as against expression in the same cell (Cell C). We hypothesized that expression of FAST and MaCAR in the same cell could directly result in TPAL, potentially fully circumventing TPA accumulation. First, we showed that supplemented TPA is not consumed under well-buffered conditions **(Fig. 3H)**. We then tested if supplying MHET or BHET would yield TPAL. Excitingly, we observed TPAL synthesis in all reaction conditions starting from MHET and BHET, indicating both strategies are viable alternatives to avoid challenges posed by TPA uptake. Using Cell C, we obtained high TPAL yields of 82.7% or 98.0% from 5 mM MHET or BHET, respectively **(Fig. 3I-J)**. Using Cell A and B in co-culture, we found that a 1:1 inoculation ratio resulted in optimum TPAL yields of 82.6% and 68.8% from MHET and BHET, respectively. To further highlight our approach, we tested both strategies in resting cells with MHET or BHET as our starting substrates. We again saw TPAL production in all our reaction conditions starting from 5 mM MHET or BHET (**Fig. S9**). We observed high TPAL yields of 77% or 88% from MHET or BHET, respectively using Cell C. These results indicate that coupling intercellular TPA synthesis to downstream enzymatic transformations can effectively enable TPA valorization using cells.

### Whole cell valorization towards diamines starting from PET intermediate deconstruction products

Having designed efficient routes from alternative PET deconstruction products to the highly reactive platform molecule TPAL, we next designed a whole cell cascade for the biosynthesis of the difunctionalized amine, *para*-xylylenediamine (pXYL). To do this, we used a transaminase from *Chromobacterium violaceum*, which has been shown to accept PET-derived aldehydes^36^. Additionally, we co-expressed an alanine dehydrogenase from *Bacillus subtilis* (AlaDH) to recycle pyruvate back to the alanine amine donor. We first investigated the operating pH range of CvTA in resting whole cells. We noticed that as reaction pH is lowered, *para*-aminomethyl benzoic acid (pAMBA) accumulates, presumably via endogenous oxidation (**Fig 4A**). We observed that the amination of 5 mM TPAL had an optimal conversion to pXYL at pH 7.5, resulting in a 90% conversion after 1 h (**Fig. 4B**). These results highlight a unique challenge if we were to start with TPA as an initial substrate as low pH is needed for TPA uptake, but higher pH is optimal for amination. Despite our progress with alternative PET deconstruction products, we chose to investigate addressing this when starting from TPA by employing a one-pot, two-step approach (**Fig. 4C**) in which the reaction is pH-adjusted from that optimal for TPA to that optimal for CvTA. For this reaction, we supplemented 5 mM TPA at a starting pH of 7.5 or 5.2. For the pH 5.2 case, after 2 h we increased pH to 7.5 with NaOH. After which, cells expressing CvTA and AlaDH (Cell D) were supplemented in both conditions and the reactions were ran for an additional 2 h. As expected, when starting at a pH of 7.5, we observed no TPA conversion (**Fig. 4D**). However, when starting at a pH of 5.2, we obtained a 97% conversion to the desired diamine pXYL (**Fig. 4E**).

**Fig. 4.**
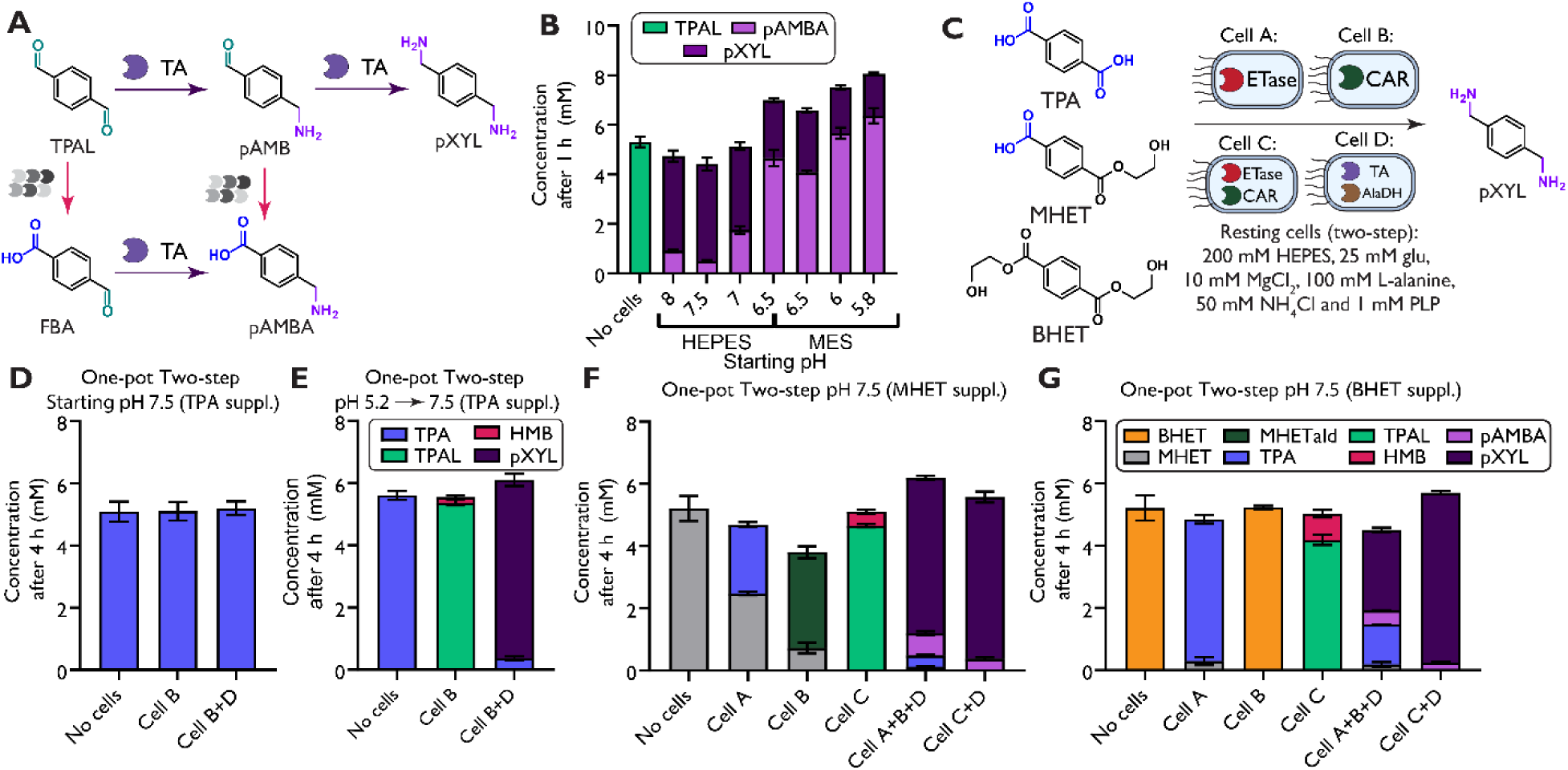
Valorization towards *para*-xylylenediamine (pXYL) using enhanced uptake of MHET and BHET. (**A**) Endogenous oxidation can limit pXYL production in resting cells. (**B**) pXYL biosynthesis from TPAL at varying pH ranges. (**C**) One-pot two-step production of pXYL from TPA, MHET, and BHET. TPA was added to reactions mixtures of Cell B at a starting pH of (**D**) 7.5 or (**E**) 5.2. After 2 h, the reaction mixture was titrated if needed to 7.5 and Cell D was added. (**F**) MHET and (**G**) BHET were added to Cell A, Cell B, Cell C as well as optimum ratio of Cell A and B found previously. After 2 h, Cell D was added. Samples sizes are *n* = 3 using biological replicates. Data shown are mean ± s.d.

While we achieved high conversion to pXYL from TPA using pH adjustment, we next aimed to produce amines without pH adjustment using MHET or BHET as starting substrates. For direct comparisons of pXYL conversion from TPA to that of our proposed cascade starting from MHET or BHET, we continued to run our reactions in a one-pot two-step approach, meaning that Cell D was added after 2 h. We ran reactions using our Cell C as well as the best performing Cell A:B ratio to generate TPAL from MHET and BHET. Using Cells A, B and D, we obtained pXYL yields of 80.5% from MHET or 57.4% from BHET **(Fig. 4 F, G**). With Cell C and D, we observed high pXYL yields of 93.1% from MHET or 95.9% from BHET. We also examined whether we could selectively biosynthesize the antifibrinolytic drug, *para*-aminomethyl benzoic acid (pAMBA) from our intermediate deconstruction products ^50^. As MaCAR has been shown to have minimal activity on pAMBA, we hypothesized that we could target pAMBA synthesis using a one-pot one-step approach^36^. We found that our three-cell approach at a cellular ration of 1:10:2 of Cells A:B:D resulting in 83% yield pAMBA from 5 mM MHET (**Fig. S10**).

### Reaction scaleup and upcycling of post-consumer PET waste

Given that coupling FAST and MaCAR in the same cell showed high TPAL and pXYL yields at 5 mM concentrations of either MHET or BHET, we investigated the ability of our upcycling cascade for either substrate at elevated substrate loading of 10, 20, or 40 mM. We ran these reactions at an increased glucose concentration (80 mM) to ensure sufficient co-factors for carboxylate reduction. For either substrate, we sampled at 4 and 8 h. We observed high yields of TPAL in just 4 h with each concentration tested for both supplemented MHET (10 mM: 77%; 20 mM: 83%; 40 mM: 80%) and BHET (10 mM: 77%; 20 mM: 79%; 40 mM: 79%) (**Fig. S11**). At 8 h, we found the yield at 40 mM increased slightly for MHET (84%) and BHET (89%).

After successfully achieving TPAL production at increased substrate loading, we scaled up our cascade to a 10 mL reaction volume (from 300 µL) and used intermediate PET deconstruction substrates from glycolysis of post-consumer textile or plastic waste (**Fig. 5A**). Using previously established methods, we performed catalytic glycolysis on a 100% polyester red T-shirt resulting in mixture of 78% BHET, 21% MHET and 1% TPA^51^. We also conducted glycolysis on a post-consumer PET plastic bottle that resulted in a mixture of 99% BHET and 1% MHET mixture^52^. We next ran our one-pot two-step method to target pXYL production starting from these deconstruction products. Both solid residues were resuspended in our cellular reaction buffer at an estimated 40 mM concentration. We supplemented Cell C for 8 h and then added Cell D and let the reaction run for an addition 8 h for a total of 16 h. We were excited to see the full conversion of deconstruction products in both reactions, with pXYL yields of 89.5% or 90.5% from polyester textile waste or post-consumer PET plastic waste, respectively (**Fig. 5B, C**). As both PET deconstruction and upcycling steps resulted in high yields, our tandem chemical deconstruction and biological upcycling strategy resulted in overall pXYL yields of >80%.

**Fig. 5.**
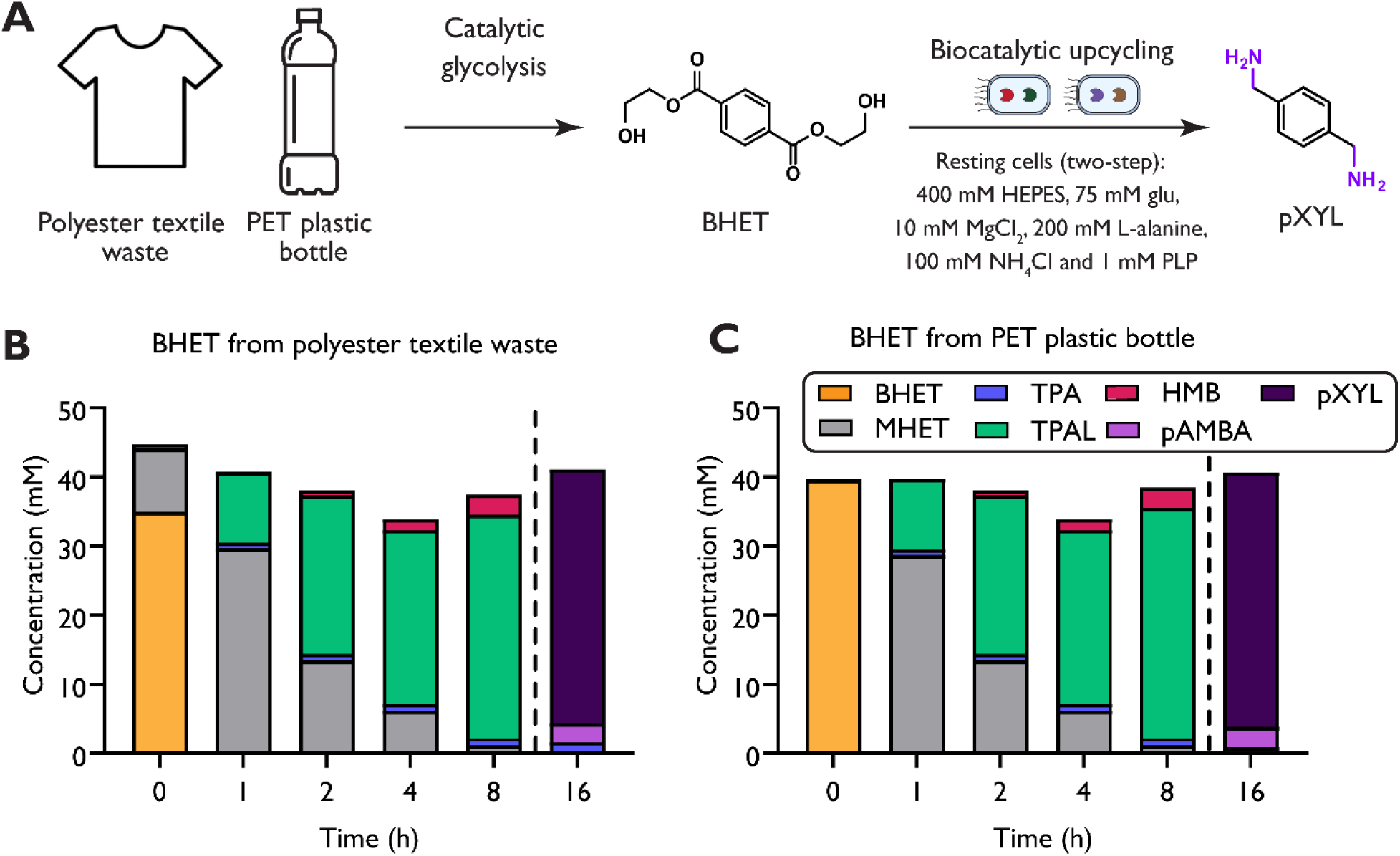
Microbial upcycling of PET deconstruction products. (**A**) Depiction of our chemoenzymatic platform for pXYL production starting from PET. Catalytic glycolysis was done on polyester textile waste as well as PET plastic bottles. These products were then coupled with the proposed whole cell cascade for the biosynthesis of pXYL. Cell C was added to resuspended PET deconstruction products from (**B**) polyester textile waste (red shirt) and (**C**) PET bottle. At 8 h, Cell D was added, and the reaction was conducted for an additional 8 h. Samples sizes are *n* = 1.

## Discussion

Prior pioneering work reporting tandem catalytic and biological processes focused on deconstruction to TPA alongside heterologous expression of TPA transporters^4,53^. Our work reinforces the value of tandem hybrid processes and provides greater reasoning for their efficient performance. Based on our findings, an entirely cellular alternative could also be designed where a secreted ETase is chosen for the ability to selectively produce glycol esters for subsequent intracellular processing by a complementary intracellular ETase. Collectively, our results showcase the industrial promise of bypassing TPA as a substrate, harnessing intracellular ETases, exploiting extensive strain engineering for TPAL retention, and interfacing engineered bacterial cells with catalytic glycolysis for robust hybrid PET valorization.

The cellular transport of various PET deconstruction products revealed in this study, if similar for other bacteria, offers new perspectives on why known natural PET degrading bacteria such as *Ideonella sakaiensis* utilize a two-enzyme (PETase-MHETase) system for PET assimilation. Naturally observed PETases tend to generate MHET, which may leverage aromatic carboxylate transporters for cellular entry that could be widespread across environmental bacteria. As a result, MHETase secretion may not be strictly necessary for natural PET assimilators. However, based on sequence conservation of the signal peptide on characterized MHETases and uncharacterized homologs, we expect MHETase secretion to be the norm. Given that, another plausible explanation is that MHETase secretion accompanied by evolution of a dedicated TPA transporter, while less efficient for the PET assimilator, may have been an evolutionary solution to mitigate the formation of MHET as a common good that other bacteria containing promiscuous hydrolases could utilize.

## Materials and methods

### Strains and plasmids

*Escherichia coli* strains and plasmids used are listed in **Table S1**. Molecular cloning and vector propagation were performed in DH5α (NEB). Polymerase chain reaction (PCR) based DNA replication was performed using KOD XTREME Hot Start Polymerase (MilliporeSigma) for plasmid backbones. Cloning was performed using Gibson Assembly. Oligos for PCR amplification and translational knockouts are shown in **Table S2**. Oligos were purchased from Integrated DNA Technologies (IDT). The DNA sequence and translated sequence of proteins overexpressed in this paper are found in **Table S3**. The pORTMAGE-Ec1 recombineering plasmid was kindly provided by Timothy Wannier and George Church of Harvard Medical School.

### Chemicals

The following compounds were purchased from MilliporeSigma: sodium borate decahydrate, sodium phosphate dibasic anhydrous, chloramphenicol, kanamycin sulfate, dimethyl sulfoxide (DMSO), boric acid, L-alanine, and HEPES. D-glucose and m-toluic acid. The following compounds were purchased from Alfa Aesar: agarose and ethanol were purchased. The following compounds were purchased from Fisher Scientific: isopropyl ß-D-1-thiogalactopyranoside (IPTG), acetonitrile, sodium chloride, trifluoroacetic acid, LB Broth powder (Lennox), and LB Agar powder (Lennox). A MOPS EZ rich defined medium kit was purchased from Teknova. Taq DNA ligase was purchased from GoldBio. Anhydrotetracycline (aTc) was purchased from Cayman Chemical. Phusion DNA polymerase and T5 exonuclease were purchased from New England BioLabs (NEB). Sybr Safe DNA gel stain was purchased from Invitrogen. The following compounds were purchased from TCI America: Pyridoxal 5’-phosphate (PLP), *ortho*-phthalaldehyde and 3-mercaptopropionic acid.

### Culture conditions

Cultures were grown in LB-Lennox medium (LB: 10 g/L bacto tryptone, 5 g/L sodium chloride, 5 g/L yeast extract) or MOPS EZ rich defined media (Teknova M2105) with 2% glucose (MOPS media). To prepare cells for resting cell assays, confluent overnight cultures of *E. coli* strains were used to inoculate 200 mL cultures in LB media in 1 L baffled shake flasks. The cultures were grown at 37°C until mid-exponential phase (OD_600_ = 0.5−0.8) and then dropped to 18°C overnight for 18 h. Cells were then pelleted and used or frozen at −80°C. Cells used in this study were stored at −80°C for less than 24 h.

### Aldehyde Stability Assays

For testing stability in metabolically active cells under aerobic growth conditions, cultures of each *E. coli* strain to be tested were inoculated from a frozen stock and grown to confluence overnight in 5 mL of LB media. Confluent overnight cultures were then used to inoculate experimental cultures to an initial starting OD_600_ of 0.01 in 400 µL volumes in a 96-deep-well plate (Thermo Scientific™ 260251) and grown at 37°C. Cultures were supplemented with 5 mM of aldehyde (prepared in 100 mM stocks in 100% DMSO) at mid-exponential phase (OD_600_ = 0.5-0.8). Cultures were then incubated at 30°C with shaking at 1000 RPM and an orbital radius of 3 mm. Samples were taken by pipetting 25 µL from the cultures into 125 µL of a 1:4 1 M HCl to methanol mixture. Samples were then centrifuged in a different 96-deep-well plate and the extracellular broth was collected. Compounds were quantified over a 24 h period using HPLC with samples collected at 4 h and 24 h.

For resting cell stability testing, cell pellets were thawed and then washed with 200 mM HEPES, pH 7.5 buffer. The mass of cell pellets was then measured, and the pellets were resuspended in 200 mM HEPES, pH 5.8 and 7.5 and at a wet cell weight of 50 mg/mL. The resuspended resting cells were then aliquoted into 96-deep-well plates and supplemented with 5 mM aldehydes of interest (prepared in 100 mM stocks in DMSO) at a reaction volume of 400 µL. Resting cells were then incubated at 30°C with shaking at 1000 RPM and an orbital radius of 3 mm. Samples were taken by pipetting 25 µL from the cultures into 125 µL of a 1:4 1 M HCl to methanol mixture. Samples were then centrifuged in a different 96-deep-well plate and the extracellular broth was collected. Compounds were quantified over a 20 h period using HPLC with samples collected at 4 h and 20 h.

### Growing cell assays

For metabolically active CAR assays, pZE plasmids expressing MaCAR and the Sfp protein from *Bacillus subtilis* were transformed into RARE.Δ16 cultures. Strains were then inoculated from a frozen stock and grown to confluence overnight in 5 mL of LB media with 50 µg/mL kanamycin. Confluent overnight cultures were then used to inoculate experimental cultures in a 96-deep-well plate initial starting OD_600_ of 0.05 in 300 µL in MOPS media with 10 mM MgCl_2_ and 50 µg/mL kanamycin at pH 7.5. For experiments testing the effect of starting pH, the starting pH of the MOPS media was titrated to pH ranging from 7.4-6.75. For experiments testing media buffering capacity, 100 mM monobasic dihydrogen phosphate were added to cultures. At mid-exponential phase, we induced each culture then supplied 5 mM acid substrate (prepared in 100 mM stocks in 100% DMSO). Cultures were then incubated at 30°C with shaking at 1000 RPM and an orbital radius of 3 mm. Samples were taken by pipetting 25 µL from the cultures into 125 µL of a 1:4 1 M HCl to methanol mixture. Samples were then centrifuged in a different 96-deep-well plate and the extracellular broth was collected. Compounds were quantified over a 24 h period using HPLC with samples collected at 4 h and 24 h. For pH tracking experiments, confluent overnight cultures were then used to inoculate 3 mL of MOPs media in 14 mL culture tubes. Cultures were supplemented with 5 mM of TPA (prepared in a 100 mM stock in DMSO) at mid exponential phase. Cultures were then incubated at 30°C in a rotor drum (Thermo Scientific Cel-Gro Tissue Culture Rotator) at maximum speed. Sampling was done as previously mentioned. Compounds were quantified and the pH was measured for 24 h with timepoints at 0 h, 4 h, 8 h, 20 h, and 24 h using HPLC.

For metabolically active ETase assays, pET-28a(+) plasmids expressing each ETase variant were transformed into RARE.Δ16. Confluent overnight cultures were then used to inoculate experimental cultures in a 96-deep-well plate initial starting OD600 of 0.05 in 300 µL in MOPS media with 10 mM MgCl2 and 50 µg/mL kanamycin at pH 7.5. At mid-exponential phase, we induced each culture then supplied 5 mM BHET, MHET or MHET-ald (prepared in 100 mM stocks in 100% DMSO). Cultures were then incubated at 30°C with shaking at 1000 RPM and an orbital radius of 3 mm. Sampling was done as previously mentioned. Compounds were quantified over a 24 h period using HPLC with samples collected at 4 h and 24 h.

For metabolically active ETase and CAR coupled assays, pACYC-FAST PETase and pZE-MaCAR plasmids were co-transformed into RARE.Δ16. Confluent overnight cultures of pET-28a(+)-FAST PETase (Cell A), pZE-MaCAR (Cell B) and pACYC-FAST PETase + pZE-MaCAR (Cell C) were then used to inoculate experimental cultures in a monoculture, expressing the dual plasmid, or a coculture, expressing FAST and MaCAR in separate strains at varying inoculation ratios. Cultures had an initial starting OD_600_ of 0.05 in 300 µL in MOPS media with 10 mM MgCl_2_ and respective antibiotics (Cell C: 17 µg/mL chloramphenicol and 25 µg/mL kanamycin, coculture and monocultures of Cell A and B: 50 µg/mL kanamycin) at pH 7.5. At mid-exponential phase, we induced each culture induced (Cell A and B: 0.1 µg/mL aTc; Cell C: 0.1 µg/mL aTc and 1 mM IPTG) then supplied 5 mM MHET or TPA (prepared in 100 mM stocks in 100% DMSO). Cultures were then incubated at 30°C. Sampling was done as previously mentioned. Compounds were quantified over a 24 h period using HPLC with samples collected at 8 h and 24 h.

### Growth rate and protein production assay

For growth rate testing of engineered strains, confluent overnight cultures were used to inoculate experimental cultures in LB or MOPS media at 100x dilution in 200 µL volumes in a Greiner clear bottom 96 well plate (Greiner 655090). Cultures were grown for 24 h in a Spectramax i3x plate reader with medium plate shaking at 37 °C and readings at 600 nm taken every 10 min to determine growth rate of each strain. For growth rate in shake flasks, confluent overnight cultures were used to inoculate experimental cultures in 50 mL of LB media at 100x dilution in 250 mL shake flasks. Cultures were grown for 9 h at 37 °C and 250 RPMs and readings at 600 nm were taken every 30 min to determine growth rate of each strain using a BioMate160 spectrophotometer.

For growth rate testing with ETase overexpression, confluent overnight cultures were used to inoculate experimental cultures in MOPS media with 50 µg/mL kanamycin at 100x dilution in 200 µL volumes in a Greiner clear bottom 96 well plate (Greiner 655090). Cultures were grown for 3 h in a Spectramax i3x plate reader with medium plate shaking at 37 °C with absorbance readings at 600 nm taken every 10 min to determine growth rate of each strain. Cultures were then induced with 1 mM IPTG and dropped to 30 °C for an additional 21 h.

### TPA protonation state calculation

The protonation state of TPA was calculated using Henderson-Hasselbalch relationship using the pK_a1_ = 3.54 and pK_a2_ = 4.46. The following equations were then used to plot the fraction of protonated species at the pH range of interest (pH 4.0-8.0) in Fig. S6:

(1) D =1+10 ^(pH − pKa1)^ +10 ^(2pH − pKa1 − pKa2)^ where D is the denominator fractional abunbances
(2) α_TPA_ = 1 / D where α_TPA_ is the fraction of fully protonated TPA
(3) α_TPA_ ^−^ = 10^(pH - pKa1)^ / D where α_TPA_ ^−^ is the fraction of partially protonated TPA
(4) α_TPA_ ^2-^ = 10^(2pH - pKa1 - pka2)^ / D where α_TPA_ ^2-^ is the fraction of fully deprotonated TPA

### Translational genomic knockouts

Translational knockouts to the RARE.Δ16 were performed using 10 rounds of multiplexed automatable genome engineering (MAGE) where stop codons were introduced into the genomic sequence for each aldehyde dehydrogenase target in the upstream portion of the gene. Construction of ROARΔ22 was done using the same oligos that were listed for the ROAR.Δ12 strain. MAGE was performed using the pORTMAGE-Ec1 recombineering plasmid. Briefly, bacterial cultures were inoculated with 1:100 dilution in 3 mL of LB media with 30 μg/mL kanamycin (Kan) and grown at 37°C until OD_600_ of 0.4-0.6 was reached. Then, the proteins responsible for recombineering were induced with 1 mM *m*-toluic acid and cultures were grown at 37°C for an additional 15 min. Cells were then prepared for electroporation by washing 1 mL three times with refrigerated 10% glycerol and then resuspending in 50 µL of 10% glycerol with each knockout oligo within a subset added at 1 µM. Cells were then electroporated and recovered in 3 mL of LB with 30 µg/mL kanamycin to be used for subsequent rounds. Preliminary assessment of knockouts was performed using mascPCR (multiplexed allele specific colony PCR) and confirmed using Sanger Sequencing. ROAR.Δ22 was cured of the pORTMAGE-Ec1 plasmid following confirmation of genomic knockouts.

### Resting whole cell assays

To prepare cells for resting cell assays, pET-28a(+)-FAST ETase (Cell A), pZE-MaCAR (Cell B), and pACYC-CvTA-AlaDH (Cell D) were transformed into ROAR.Δ22. pACYC-FAST ETase and pZE-MaCAR (Cell C) were also co-transformed into ROAR.Δ22. Confluent overnight cultures were used to inoculate 200 mL cultures in LB media antibiotics (Cell A and B: 50 µg/mL kanamycin; Cell C: 25 µg/mL kanamycin and 17 µg/mL chloramphenicol; Cell D: 34 µg/mL chloramphenicol) with 1 L baffled shake flasks. The cultures were grown at 37°C until mid-exponential phase and then induced (Cell A and B: 0.1 µg/mL aTc; Cell C: 0.1 µg/mL aTc and 1 mM IPTG; Cell D: 1 mM IPTG). After induction the temperature was dropped to 18°C overnight for 18 h. Cells were then pelleted, and the mass of the cell pellets were then measured. Cells were used immediately after overexpression or frozen at −80°C. Cells used in this study were stored at −80°C for less than 24 h. All cells were washed with 200 mM HEPES, pH 7.5 buffer prior to use. Reactions unless otherwise noted were performed at a reaction volume of 400 µL in 1 mL deep-well plates.

To assay MaCAR (Cell B) in resting cells, cells were resuspended in buffer with 200 mM HEPES or 200 mM MES (dependent on starting pH), 25 mM glucose, 10 mM magnesium chloride at pH 5.2-7.5. The resuspended resting cells were then aliquoted into 96-deep-well plates at a wet cell weight of 50 mg/mL and supplemented with 5 mM substrate of interest (prepared in 100 mM stocks in DMSO) at a reaction volume of 400 µL. Resting cells were then incubated at 30°C with shaking at 1000 RPM and an orbital radius of 3 mm. Samples were taken by pipetting 25 µL from the cultures into 125 µL of a 1:4 1 M HCl to methanol mixture. Samples were then centrifuged in a different 96-deep-well plate and the extracellular broth was collected. Compounds were quantified over a 4 h period using HPLC with samples collected at 2 h and 4 h.

To assay FAST ETase and MaCAR (Cell A+B and Cell C) in resting cells, cells were resuspended in buffer with 200 mM HEPES, 25 mM glucose, 10 mM magnesium chloride at a pH 7.5. The resuspended resting cells were then aliquoted to a total wet cell weight of 50 mg/mL at varying ratios of FAST ETase to MaCAR cells. Reaction mixture was supplemented with 5 mM BHET, MHET or TPA at a volume of 400 µL. Resting cells were then incubated at 30°C. Sampling was done was previously mentioned. Compounds were quantified over a 4 h period using HPLC with samples collected at 2 h and 4 h.

To assay Cell C with high substrate loading, cells were resuspended in buffer with 400 mM HEPES, 75 mM glucose, 10 mM MgCl_2_ at pH 7.5. Cells were supplemented with 10, 20, or 40 mM of BHET or MHET at a cell concentration of 50 mg_WCW_/mL and at a volume of 400 µL. Compounds were quantified over an 8 h period using HPLC with samples collected at 4 h and 8 h.

To assay CvTA-AlaDH (Cell D) in resting cells, cells were resuspended in buffer with 200 mM HEPES or 200 mM MES, 25 mM glucose, 10 mM MgCl2, 100 mM L-alanine, 50 mM NH_4_Cl and 1 mM PLP at pH 5.8-8.0. The resuspended resting cells were then aliquoted to a total wet cell weight of 50 mg/mL and supplemented with 5 mM TPAL. Resting cells were then incubated at 30°C. Sampling was done was previously mentioned. Compounds were quantified using HPLC at 2 h.

To assay Cell B and D in a one-pot two-step method, cells were resuspended in buffer with 200 mM HEPES, 200 mM MES, 25 mM glucose, 10 mM MgCl_2_, 100 mM L-alanine, 50 mM NH_4_Cl and 1 mM PLP at a pH of 5.2 or 7.5. To start the reaction, Cell B was added to the reaction mixture at 50 mg_WCW_/mL and at a starting pH of 5.2 or 7.5. The cells were then supplemented with 5 mM TPA disodium salt and incubated at 30°C. After 2 h, the low starting pH reaction mixtures were treated to 7.5 using NaOH. Also, at 2 h both starting pH reactions were supplemented with 50 mg_WCW_/mL of CvTA cells. The reactions were then ran for an additional 2 h. Sampling was done was previously mentioned. Compounds were quantified at 4 h using HPLC.

To assay Cell A+B, Cell C and Cell D in one-pot with resting cells, cells were resuspended in buffer with 200 mM HEPES, 25 mM glucose, 10 mM MgCl_2_, 100 mM L-alanine, 50 mM NH_4_Cl and 1 mM PLP at a pH of 7.5. For pXYL targeted reactions, Cells A and B were added to the reaction mixture at different ratios (1:2, 1:1. 2:1, 1:5 and 1: 10 of Cell A: Cell B) with a total cell concentration of 50 mg_WCW_/mL. Cell C was also added at a cell concentration of 50 mg_WCW_/mL. The cells were then supplemented with 5 mM BHET, MHET or TPA and incubated at 30°C. After 2 h, 50 mg_WCW_/mL of Cell D was added and the reaction was then ran for an additional 2 h. Sampling was done was previously mentioned. Compounds were quantified at 4 h using HPLC. For pAMBA targeted reactions, Cells A, B and D were added to the reaction mixture at different ratios (2:1:1, 1:2:1. 1:10:2, and 1:1:2 of Cell A: Cell B: Cell D) with a total cell concentration of 50 mg_WCW_/mL. Cells C and D were also added at different ratios (10:1, 5:1 and 1:1 of Cell C: CellD) with a total cell concentration of 50 mg_WCW_/mL. The cells were then supplemented with 5 mM MHET and incubated at 30°C. Sampling was done was previously mentioned. Compounds were quantified at 4 h using HPLC.

### Amine production from real PET glycolysis products

To assay Cell C using real PET glycolysis deconstruction products, cells were resuspended in buffer with 400 mM HEPES, 75 mM glucose, 10 mM MgCl_2_, 200 mM L-alanine, 100 mM NH_4_Cl and 1 mM PLP at pH 7.8. Solid PET glycolysis products were resuspended in 500 µL DMSO and added to the reaction mixture at a targeted 40 mM BHET concentration. Reactions were conducted using a 50 mL conical tube with a total reaction volume of 10 mL. Sampling was done was previously mentioned. Compounds were quantified over an 8 h period using HPLC with samples collected at 4 h and 8 h.

### PET glycolysis

Plastic PET glycolysis experiments were carried out under previously optimized conditions^52^ using a Monowave 450 microwave reactor (Anton Paar GmbH), equipped with an in-built IR sensor and an external Ruby thermometer for precise temperature control, Commercial PET beverage bottles (Coca-Cola) were washed with water, dried, and cut into ∼4 mm × 4 mm square pieces prior to use.

In a typical experiment, 500 mg of PET flakes, 5 mL of ethylene glycol (EG), and 5 mg of catalyst were added to a microwave reaction vial. Commercial ZnO nanopowder (<5 μm particle size, Sigma-Aldrich) was used as the catalyst. The contents of the vial were thoroughly mixed using a vortex mixer to ensure uniform dispersion of ZnO in EG. The vial was then placed in the microwave reactor, which was programmed to maintain a constant temperature of 210°C for 15 minutes with a stirring speed of 600 rpm.

After the reaction, the vial was rapidly cooled to room temperature. Subsequently, 100 mL of distilled water was added to extract BHET. Unreacted PET and higher molecular weight oligomers were removed by filtration using Whatman filter paper. The filtrate, containing water-soluble products, was analysed by high-performance liquid chromatography (HPLC).

The residual water in the product solution was evaporated under vacuum (72 mbar) at 40°C using a rotary evaporator. BHET was then crystallized by cooling the concentrated solution overnight at 4°C in a refrigerator, following the addition of a small volume of distilled water. The resulting BHET crystals were collected by filtration using a glass filter, thoroughly washed to remove any residual EG, and dried overnight at 80°C.

The conversion of PET is calculated as follows:

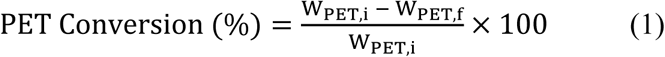

W_PET,i_ corresponds to the initial weight of PET and W_PET,f_ to the weight of the unreacted PET, obtained via filtration. The yield of the BHET is defined as:

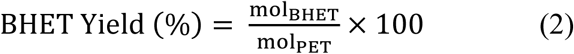

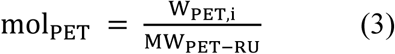

Textile depolymerization experiments were carried out at previously optimized conditions^51^. Depolymerization experiments were performed on a Monowave 450 microwave reactor (Anton Paar GmbH). This batch microwave reactor controls the temperature, time, and maximum set power. Typically, 500 mg of textiles, 5 mL of EG, and 5 mg of the catalyst were placed in a microwave reaction vial. The vial was inserted into the microwave reactor, and the reactor was programmed to maintain a constant temperature. On completion of a reaction, the reaction vial was allowed to cool down rapidly to room temperature. 100 mL of distilled water was added to separate BHET and oligomers. The unreacted polymer and larger oligomers were removed using a Whatman filter paper. The residual water from the product solution was then evaporated under vacuum (72 mbar) at 60 °C using a rotary evaporator, and the resulting BHET was crystallized by cooling the residual solution overnight to 4 °C in a refrigerator after the addition of a small amount of distilled water to the residue. The resultant crystals were filtered using a glass filter and dried at 80 °C.

### HPLC analysis

TPAL, 4HMB, 4HMBA, BDM and pXYL were quantified using reverse-phase high-performance liquid chromatography (HPLC) with an Agilent 1260 Infinity with a Zorbax Eclipse Plus-C18 column with a guard column installed. Amines were derivatized with *ortho*-pthalaldehyde and 3-mercaptopropionic acid and identified using reverse-phase high-performance liquid chromatography (RP-HPLC) with an Agilent 1260 Infinity with a Zorbax Eclipse Plus-C18 column with a guard column installed. The methods for detection of TPAL, 4HMB, 4HMBA, BDM and pXYL were described previously^36^.

## Acknowledgements

The authors would like to thank the members of the University of Delaware Center for Plastics Innovation for guidance and support on this project.

## Author Contributions

Conceptualization: RMD, AMK

Data curation: RMD, ES, EA

Methodology: RMD, ES, EA

Investigation: RMD, ES, EA, PN

Visualization: RMD

Funding acquisition: AMK

Project administration: AMK, DGV

Supervision: AMK, DGV

Writing – original draft: RMD, AMK

Writing – review & editing: RMD, AMK, ES, EA, DGV

## Data and materials availability

All data are available in the main text or the supplementary materials. Strains and plasmids are available upon request.

## Supplementary Materials

Tables S1 to S4

Figs. S1 to S10

Supplementary references

## References

1. Pottinger, A. S. et al. Pathways to reduce global plastic waste mismanagement and greenhouse gas emissions by 2050. Science 386, 1168–1173 (2024).

2. Liu, F. et al. Current advances in the structural biology and molecular engineering of PETase. Front. Bioeng. Biotechnol. 11, (2023).

3. Taek Kim, H., et al. Biological Valorization of Poly(ethylene terephthalate) Monomers for Upcycling Waste PET. ACS Sustain. Chem. Amp Eng. 7, 19396–19406 (2019).

4. Werner, A. Z. et al. Tandem chemical deconstruction and biological upcycling of poly(ethylene terephthalate) to β-ketoadipic acid by *Pseudomonas putida* KT2440. Metab. Eng. 67, 250–261 (2021).

5. Diao, J., Hu, Y., Tian, Y., Carr, R. & Moon, T. S. Upcycling of poly(ethylene terephthalate) to produce high-value bio-products. Cell Rep. 42, (2023).

6. Dissanayake, L. & Jayakody, L. N. Engineering Microbes to Bio-Upcycle Polyethylene Terephthalate. Front. Bioeng. Biotechnol. 9, (2021).

7. Johnson, N. W. et al. A biocompatible Lossen rearrangement in Escherichia coli. Nat. Chem. 17, 1020–1026 (2025).

8. Gopal, M. R. & Kunjapur, A. M. Harnessing biocatalysis to achieve selective functional group interconversion of monomers. Curr. Opin. Biotechnol. 86, 103093 (2024).

9. Vermaas, J. V. et al. Passive membrane transport of lignin-related compounds. Proc. Natl. Acad. Sci. 116, 23117–23123 (2019).

10. Li, T. & Crook, N. Construction of a Heterologous Pathway in Escherichia coli for Terephthalate Assimilation. 2025.02.25.640003 Preprint at 10.1101/2025.02.25.640003 (2025).

11. Li, Y. et al. Metabolic engineering of *Escherichia coli* for upcycling of polyethylene terephthalate waste to vanillin. Sci. Total Environ. 957, 177544 (2024).

12. Sadler, J. C. & Wallace, S. Microbial synthesis of vanillin from waste poly(ethylene terephthalate). Green Chem. 23, 4665–4672 (2021).

13. Valenzuela-Ortega, M., Suitor, J. T., White, M. F. M., Hinchcliffe, T. & Wallace, S. Microbial Upcycling of Waste PET to Adipic Acid. ACS Cent. Sci. 9, 2057–2063 (2023).

14. Wang, S. et al. Recyclable, Malleable, and Strong Thermosets Enabled by Knoevenagel Adducts. J. Am. Chem. Soc. 146, 9920–9927 (2024).

15. Granado, L., Tavernier, R., Foyer, G., David, G. & Caillol, S. Catalysis for highly thermostable phenol-terephthalaldehyde polymer networks. Chem. Eng. J. 379, 122237 (2020).

16. Granado, L. et al. Toward Sustainable Phenolic Thermosets with High Thermal Performances. ACS Sustain. Chem. Eng. 7, 7209–7217 (2019).

17. Sonmez, H. B., Kuloglu, F. G., Karadag, K. & Wudl, F. Terephthalaldehyde- and isophthalaldehyde-based polyspiroacetals. Polym. J. 44, 217–223 (2012).

18. Katsoulidis, A. P. et al. Copolymerization of terephthalaldehyde with pyrrole, indole and carbazole gives microporous POFs functionalized with unpaired electrons. J. Mater. Chem. A 1, 10465–10473 (2013).

19. Li, Z. et al. Removal and adsorption mechanism of tetracycline and cefotaxime contaminants in water by NiFe2O4-COF-chitosan-terephthalaldehyde nanocomposites film. Chem. Eng. J. 382, 123008 (2020).

20. Brindell, G. D., Lillwitz, L. D., Wuskell, J. P. & Dunlop, A. P. Polymer Applications of Some Terephthalaldehyde Derivatives. Prod. RD 15, 83–88 (1976).

21. Chung, Y., Hyun, K. H. & Kwon, Y. Fabrication of a biofuel cell improved by the π-conjugated electron pathway effect induced from a new enzyme catalyst employing terephthalaldehyde. Nanoscale 8, 1161–1168 (2015).

22. Díez, N., Mysyk, R., Zhang, W., Goikolea, E. & Carriazo, D. One-pot synthesis of highly activated carbons from melamine and terephthalaldehyde as electrodes for high energy aqueous supercapacitors. J. Mater. Chem. A 5, 14619–14629 (2017).

23. Choi, S.-H., Yashima, E. & Okamoto, Y. Synthesis of Polyesters from Terephthalaldehyde and Isophthalaldehyde through Tishchenko Reaction Catalyzed by the Ethylmagnesium Bromide-(−)-Sparteine Complex and Aluminum Alkoxides. Polym. J. 29, 261–268 (1997).

24. Weng, Q. et al. Controllable Synthesis and Biological Application of Schiff Bases from d-Glucosamine and Terephthalaldehyde. ACS Omega 5, 24864–24870 (2020).

25. Shaygan, S., Pasdar, H., Foroughifar, N., Davallo, M. & Motiee, F. Cobalt (II) Complexes with Schiff Base Ligands Derived from Terephthalaldehyde and ortho-Substituted Anilines: Synthesis, Characterization and Antibacterial Activity. Appl. Sci. 8, 385 (2018).

26. Gavali, L. V. et al. Novel terephthalaldehyde bis(thiosemicarbazone) Schiff base ligand and its transition metal complexes as antibacterial Agents: Synthesis, characterization and biological investigations. Results Chem. 7, 101316 (2024).

27. Kunjapur, A. & Prather, K. L. Microbial Engineering for Aldehyde Synthesis. Appl. Environ. Microbiol. 81, 1892–1901 (2015).

28. Zhou, J., Chen, Z. & Wang, Y. Bioaldehydes and beyond: Expanding the realm of bioderived chemicals using biogenic aldehydes as platforms. Curr. Opin. Chem. Biol. 59, 37–46 (2020).

29. Schober, L. et al. Enzymatic reactions towards aldehydes: An overview. Flavour Fragr. J. 38, 221–242 (2023).

30. Zhubanov, B. A., Kravtsova, V. D., Zhubanov, K. A., Abildin, T. S. & Bizhanova, N. B. p-Xylylenediamine and Its New Polyimides. Eurasian Chem.-Technol. J. 6, 45–50 (2004).

31. Stavila, E., Alberda van Ekenstein, G. O. R. & Loos, K. Enzyme-Catalyzed Synthesis of Aliphatic–Aromatic Oligoamides. Biomacromolecules 14, 1600–1606 (2013).

32. Nanclares, J., Petrović, Z. S., Javni, I., Ionescu, M. & Jaramillo, F. Segmented polyurethane elastomers by nonisocyanate route. J. Appl. Polym. Sci. 132, (2015).

33. Dataintelo. P xylylenediamine Market Report | Global Forecast From 2025 To 2033. https://dataintelo.com/report/global-p-xylylenediamine-market.

34. Report, F. M. P-xylylenediamineMarket Size, Share, Growth | CAGR Forecast2032. Future Market Report https://www.futuremarketreport.com/industry-report/p-xylylenediamine-market/.

35. P Xylylenediamine Market Size, Growth, Share, & Analysis Report - 2033. https://datahorizzonresearch.com/p-xylylenediamine-market-43216.

36. Gopal, M. R. et al. Reductive Enzyme Cascades for Valorization of Polyethylene Terephthalate Deconstruction Products. ACS Catal. 4778–4789 (2023) doi:10.1021/acscatal.2c06219.

37. Dickey, R. M., Jones, M. A., Butler, N. D., Govil, I. & Kunjapur, A. M. Genome engineering allows selective conversions of terephthalaldehyde to multiple valorized products in bacterial cells. AIChE J. 69, e18230 (2023).

38. Butler, N. D., Anderson, S. R., Dickey, R. M., Nain, P. & Kunjapur, A. M. Combinatorial gene inactivation of aldehyde dehydrogenases mitigates aldehyde oxidation catalyzed by E. coli resting cells. Metab. Eng. (2023) 10.1016/j.ymben.2023.04.014.

39. Britton, D. et al. Protein-engineered leaf and branch compost cutinase variants using computational screening and IsPETase homology. Catal. Today 433, 114659 (2024).

40. Fritzsche, S., Hübner, H., Oldiges, M. & Castiglione, K. Comparative evaluation of the extracellular production of a polyethylene terephthalate degrading cutinase by Corynebacterium glutamicum and leaky Escherichia coli in batch and fed-batch processes. Microb. Cell Factories 23, 274 (2024).

41. Lu, H. et al. Machine learning-aided engineering of hydrolases for PET depolymerization. Nature 604, 662–667 (2022).

42. Bell, E. L. et al. Directed evolution of an efficient and thermostable PET depolymerase. Nat. Catal. 5, 673–681 (2022).

43. Cui, Y. et al. Computational Redesign of a PETase for Plastic Biodegradation under Ambient Condition by the GRAPE Strategy. ACS Catal. 11, 1340–1350 (2021).

44. Son, H. F. et al. Rational Protein Engineering of Thermo-Stable PETase from Ideonella sakaiensis for Highly Efficient PET Degradation. (2019) doi:10.1021/acscatal.9b00568.

45. Zhong-Johnson, E. Z. L., Voigt, C. A. & Sinskey, A. J. An absorbance method for analysis of enzymatic degradation kinetics of poly(ethylene terephthalate) films. Sci. Rep. 11, 928 (2021).

46. Tournier, V. et al. An engineered PET depolymerase to break down and recycle plastic bottles. Nature 580, 216–219 (2020).

47. Xi, X. et al. Secretory expression in *Bacillus subtilis* and biochemical characterization of a highly thermostable polyethylene terephthalate hydrolase from bacterium HR29. Enzyme Microb. Technol. 143, 109715 (2021).

48. Cui, Y. et al. Computational redesign of a hydrolase for nearly complete PET depolymerization at industrially relevant high-solids loading. Nat. Commun. 15, 1417 (2024).

49. Pfaff, L. et al. Multiple Substrate Binding Mode-Guided Engineering of a Thermophilic PET Hydrolase | ACS Catalysis. ACS Catal. 12, 9790–9800 (2022).

50. Banach, L. et al. Synthesis and Hemostatic Activity of New Amide Derivatives. Molecules 27, 2271 (2022).

51. Andini, E., Bhalode, P., Gantert, E., Sadula, S. & Vlachos, D. G. Chemical recycling of mixed textile waste. Sci. Adv. 10, eado6827 (2024).

52. Selvam, E., Luo, Y., Ierapetritou, M., Lobo, R. F. & Vlachos, D. G. Microwave-assisted depolymerization of PET over heterogeneous catalysts. Catal. Today 418, 114124 (2023).

53. Sullivan, K. P. et al. Mixed plastics waste valorization through tandem chemical oxidation and biological funneling. Science 378, 207–211 (2022).

