## Supplemental Information for "Robust cellular transformations of PET deconstruction products by import of glycol esters"

**The PDF file includes:**

Tables S1 to S4
Figs. S1 to S10
References

### Supplementary Tables

### Table S1.

Strains and plasmids used in this study.

| Name | Relevant genotype | Source |
|---|---|---|
| | ***E. coli* strains** | |
| DH5α | F– Φ80*lac*ZΔM15 Δ(*lac*ZYA-argF) U169 *rec*A1 *end*A1 *hsd*R17 (rK–, mK+) *pho*A *sup*E44 λ– *thi*-1 *gyr*A96 *rel*A1 | NEB |
| MG1655 | F- λ- *ilv*G- *rfb-50 rph-1* | ATCC 700926 |
| MG1655 (DE3) | F- λ- *ilv*G- *rfb-50 rph-1* (λ DE3)<br>λ DE3 = λ sBamHIo ΔEcoRI-B int::(*lac*I::PlacUV5::T7 gene1) i21 Δ*nin*5 | Previous study |
| RARE.Δ6 | MG1655(DE3) Δ*dkgB* Δ*yeaE* Δ*(yqhC-dkgA)* Δ*yahK* Δ*yjgB* | Previous study[4] |
| ROAR.Δ12 | RARE.Δ6 Δ*aldB* Δ*puuC* Δ*betB* Δ*patD* Δ*feaB* Δ*gabD* | Previous study[5] |
| RARE.Δ16 | RARE.Δ6 Δ*adhP,* Δ*fucO,* Δ*eutG,* Δ*yiaY,* Δ*adhE,* Δ*eutE,* Δ*gldA,* Δ*gpr,* Δ*ybbO,* Δ*yghA* | Previous study[6] |
| RARE.Δ16-port | RARE.Δ16 harboring pORTMAGE-Ec1 | Previous study[6] |
| ROAR.Δ22 | RARE.Δ16 Δ*aldB* Δ*puuC* Δ*betB* Δ*patD* Δ*feaB* Δ*gabD* | This study |
| RMD001 (Cell B) | ROAR.Δ22 harboring pZE-MAV-sfp | This study |
| RMD002 | RARE.Δ16 harboring pZE-MAV-sfp | This study |
| RMD003 | RARE.Δ16 harboring pET28a(+)-TS PETase | This study |
| RMD004 | RARE.Δ16 harboring pET28a(+)-Thermo PETase | This study |
| RMD005 (Cell A) | RARE.Δ16 harboring pET28a(+)-FAST PETase | This study |
| RMD006 | RARE.Δ16 harboring pET28a(+)-PES PETase | This study |
| RMD007 | RARE.Δ16 harboring pET28a(+)-Dura PETase | This study |
| RMD008 | RARE.Δ16 harboring pET28a(+)-BHR PETase | This study |
| RMD009 | RARE.Δ16 harboring pET28a(+)-Turbo PETase | This study |
| RMD010 | RARE.Δ16 harboring pET28a(+)-LCC PETase | This study |
| RMD011 | RARE.Δ16 harboring pET28a(+)-ICCG PETase | This study |
| RMD012 | RARE.Δ16 harboring pET28a(+)-HOT PETase | This study |
| RMD013 | ROAR.Δ22 harboring pET28a(+)-FAST PETase | This study |
| RMD014 (Cell C) | ROAR.Δ22 harboring pZE-MAV-sfp and pACYC-FAST PETase | This study |
| RMD015 (Cell D) | ROAR.Δ22 harboring pACYC-CvTA-AlaDH | This study |
| | **Plasmids** | |
| pORTMAGE-EC1 | RSF1010 ori, $Kan^R$, harboring CspRecT and *mutL* (E32K) genes | Previous study |
| pZE-MAV-sfp | ColE1 ori, KanR, TetR, Tet promoter with a codon optimized CAR gene from *Mycobacterium avium* bearing an N-terminal hexahistidine tag and sfp gene from *Bacillus subtilis* | Previous study[7] |
| pET28a(+)-TS PETase | ColE1/f1 ori, KanR, LacI, T7 promoter with a codon optimized TS PETase gene bearing an C-terminal hexahistidine tag | This study |
| pET28a(+)-Thermo PETase | ColE1/f1 ori, KanR, LacI, T7 promoter with a codon optimized Thermo PETase gene bearing an C-terminal hexahistidine tag | This study |
| pET28a(+)-FAST PETase | ColE1/f1 ori, KanR, LacI, T7 promoter with a codon optimized FAST PETase gene bearing an C-terminal hexahistidine tag | This study |
| pET28a(+)-PES PETase | ColE1/f1 ori, KanR, LacI, T7 promoter with a codon optimized PES PETase gene bearing an C-terminal hexahistidine tag | This study |
| pET28a(+)-Dura PETase | ColE1/f1 ori, KanR, LacI, T7 promoter with a codon optimized Dura PETase gene bearing an C-terminal hexahistidine tag | This study |
| pET28a(+)-BHR PETase | ColE1/f1 ori, KanR, LacI, T7 promoter with a codon optimized BHR PETase gene bearing an C-terminal hexahistidine tag | This study |
| pET28a(+)-Turbo PETase | ColE1/f1 ori, KanR, LacI, T7 promoter with a codon optimized Turbo PETase gene bearing an C-terminal hexahistidine tag | This study |
| pET28a(+)-LCC PETase | ColE1/f1 ori, KanR, LacI, T7 promoter with a codon optimized LCC PETase gene bearing an C-terminal hexahistidine tag | This study |
| pET28a(+)-ICCG PETase | ColE1/f1 ori, KanR, LacI, T7 promoter with a codon optimized ICCG PETase gene bearing an C-terminal hexahistidine tag | This study |

| | | |
|---|---|---|
| pET28a(+)-HOT PETase | ColE1/f1 ori, KanR, LacI, T7 promoter with a codon optimized HOT PETase gene bearing an C-terminal hexahistidine tag | This study |
| pACYC-FAST PETase | P15a ori, $Cm^R$, LacI, Lac promoter with the codon-optimized FAST PETase gene bearing an C-terminal hexahistidine tag | This study |
| pACYC-CvTA-AlaDH (Addgene ID: 206430) | P15a ori, $Cm^R$, LacI, Lac promoter with the codon-optimized gene for ω-transaminase from *Chromobacterium violaceum* and a separate Lac promoter with the codon-optimized gene for alanine dehydrogenase from *Bacillus subtilis* | Previous study[6] |

**Table S2.**
Oligonucleotides used in this study. For MAGE oligos, the codon mutations are indicated as (Original codon_Codon position_New codon).

| Oligo Name | Sequence (5' to 3') |
|---|---|
| pET28a(+)-bb-fwd | gatccggctgctaac |
| pET28a(+)-bb-rev | ggtatatctccttcttaaagttaaac |
| pET28a(+)-TS-fwd | ctttaagaaggagatataccatgCAGACCAATCCATACG |
| pET28a(+)-Thermo-fwd | ctttaagaaggagatataccatgCAGACCAATCCATACG |
| pET28a(+)-FAST-fwd | ctttaagaaggagatataccatgCAGACCAATCCATACG |
| pET28a(+)-PES-fwd | ctttaagaaggagatataccATGGCGAACCCGTACG |
| pET28a(+)-Dura-fwd | ctttaagaaggagatataccatgCAGACCAATCCATACG |
| pET28a(+)-BHR PETase | ctttaagaaggagatataccatgAGCAATCCGTATCAGC |
| pET28a(+)-turbo-PETase | ctttaagaaggagatataccatgAGCAATCCGTATCAGC |
| pET28a(+)-LCC-PETase | ctttaagaaggagatataccatgAGCAACCCGTACCAG |
| pET28a(+)-ICCG-PETase | ctttaagaaggagatataccatgAGCAACCCGTACCAG |
| pET28a(+)-HOT PETase | ctttaagaaggagatataccATGCAGACTAACCCCTATGC |
| pET28a(+)-his-rev | gctttgttagcagccggatcTCAGTGGTGGTGGTGG |
| pACYC-FAST-fwd | ctttaataaggagatataccatgCAGACCAATCCATACG |
| pACYC-FAST-rev | gGGGATCCCCCATCAAGCTTTCAGTGGTGGTGGTGG |
| pACYC-FAST-bb-fwd | AAGCTTGATGGGGGATC |
| pACYC-FAST-bb-rev | ggtatatctccttattaaagttaaacaa |
| feaB KO oligo (TTA_9_TAA, AGC_10_TGA) | A*A*TAATAAGGAAAAGTGATGACAGAGCCGCATGTAGCAGTATAATGACAGGTCCAACAGTTTCTCGATC GTCAACACGGTCTTTATATTG |
| puuC KO oligo (TAC_8_TAA, TGG_9_TGA) | G*A*CGTGAAACAGGAGTCATAATGAATTTTCATCATCTGGCTTAATGACAGGATAAAGCGTTAAGTCTCG CCATTGAAAACCGCTTATTTA |
| betB KO oligo (GCA_5_TGA, GAA_6_TAA) | T*C*TACCCACCGATTAACCGAGGAGACGTGATGTCCCGAATGTGATAACAGCAGCTTTATATACATGGTG GTTATACCTCCGCCACCAGCG |
| gabD KO oligo (GAA_18_TAA, TGG_19_TGA) | T*T*AACGACAGTAACTTATTCCGCCAGCAGGCGTTGATTAACTGGTAATGACTGGACGCCAACAATGGTG AAGCCATCGACGTCACCAATC |
| patD KO oligo (CAT_3_TAA, TTA_5_TGA) | A*C*AATGGTCAATAACCACTGATACAGGAATATGCTATGCAATAAAAGTGACTGATTAACGGAGAACTGG TTAGCGGCGAAGGGGAAAAAC |
| aldB KO oligo (TTA_21_TGA, AAA_22_TAA) | G*G*GCTACCCATTCGCCGCCAATAAAGTTGTCATAGCGGGCTTATCACTTGAGGGGGAAACCATACTCG CCGGGCTTAATCTGTGCTGAAG |
| feaB FWD SEQ | CTTTTCTTGTCGCTGCGTACACTG |
| feaB REV SEQ | GTTCCATGGCACAATTCCCGC |
| feaB WT FWD | GACAGAGCCGCATGTAGCAGTATT |
| feaB MUT FWD | GACAGAGCCGCATGTAGCAGTATA |

| | |
|---|---|
| feaB REV | GGCAGACATGACTGCGTTATCTACATC |
| puuC FWD SEQ | AGAGGCGGCTTGAAGGATGAAG |
| puuC REV SEQ | GTTCAAATACGCCGCGTGC |
| puuC WT FWD | AGGAGTCATAATGAATTTTCATCATCTGGCTTAC |
| puuC MUT FWD | AGGAGTCATAATGAATTTTCATCATCTGGCTTAA |
| puuC REV | CCAGCTCATGGCTACTGGTGG |
| betB FWD SEQ | TCTGAGCGGCAAACCGCTG |
| betB REV SEQ | CGCGATCGACATCCTCGC |
| betB WT FWD | GGAGACGTGATGTCCCGAATGG |
| betB MUT FWD | GGAGACGTGATGTCCCGAATGT |
| betB REV | GGGCGCTTTTCACGGCG |
| gabD FWD SEQ | CGGCACGTAGTGTGGATGCC |
| gabD WT FWD | GCCAGCAGGCGTTGATTAACG |
| gabD MUT FWD | GCCAGCAGGCGTTGATTAACT |
| gabD REV | GGCACGCTACCCAGCTTGT |
| patD FWD SEQ | GTAACAACCTTGCCGATCCTGGG |
| patD WT FWD | CCACTGATACAGGAATATGCTATGCAAC |
| patD MUT FWD | CCACTGATACAGGAATATGCTATGCAAT |
| patD REV | GGTTTGCCACAATTACGGGACTCCAGT |
| aldB FWD SEQ | TTCAACTATCTCTGTAACCCTTGC |
| aldB REV SEQ | GCAATCTTAAACAGAATCGCCG |
| aldB WT FWD | GCCAATAAAGTTGTCATAGCGGGCTTT |
| aldB MUT FWD | GCCAATAAAGTTGTCATAGCGGGCTTA |
| aldB REV | TACAACCTGCCACACGGTGTATG |

**Table S3.**
Sequences of proteins expressed in this paper.

| Plasmid Name | DNA CDS | Protein Sequence |
|---|---|---|
| pZE-Ub-GFP | ATGCAGATTTTTGTGAAGACTTTAACAGGTAAGACGATTACCCTGG<br>AGGTGGAGTCCTCGGACACCATCGATAATGTAAAATCAAAAATCC<br>AAGATAAGGAAGGAATCCCTCCAGACCAGCAACGTCTGATTTTCGC<br>AGGTAAACAACTGGAGGATGGTCGCACGCTTTCGGACTACAACATC<br>CAGAAAGAATCTACCCTTCATTTGGTTCTGCGTCTGCGTGGAGGAT<br>AGTTGTTTGTGCAGGAGCTTGCATCCAAGGGCGAGGAGCTCTTTAC<br>TGGCGTAGTACCAATTCTCGTAGAGCTCGATGGCGATGTAAATGGC<br>CATAAGTTTTCCGTACGCGGCGAGGGCGAGGGCGATGCAACTAAC<br>GGCAAGCTCACTCTCAAGTTTATTTGTACTACTGGCAAGCTCCCAG<br>TACCATGGCCAACTCTCGTAACTACTCTGACCTATGGCGTACAATG<br>TTTTTCCCGCTATCCAGATCACATGAAGCAACATGATTTTTTTAAGT<br>CCGCAATGCCAGAGGGCTATGTACAAGAGCGCACTATTAGCTTTAA<br>GGATGATGGCACCTATAAGACTCGCGCAGAGGTAAAGTTTGAGGG<br>CGATACTCTCGTAAATCGCATTGAGCTCAAGGGCATTGATTTTAAG<br>GAGGATGGCAATATTCTCGGCCATAAGCTGGAGTATAATTTCAATT<br>CCCATAATGTATACATTACCGCAGATAAGCAAAAGAATGGCATTAA<br>GGCGAATTTTAAGATTCGCCATAATGTGGAGGATGGCTCCGTACAA<br>CTCGCAGATCATTATCAACAAAATACTCCAATTGGCGATGGCCCAG<br>TACTCCTCCCAGATAATCATTATCTCTCCACTCAATCCGTGCTCTCC<br>AAAGATCCAAATGAGAAGCGCGATCACATGGTACTCCTGGAGTTTG<br>TAACTGCAGCAGGCATTACTCATGGCATGGATGAGCTCTATAAGCT<br>CGAGCACCACCACCACCACCACCACTAA | MQIFVKTLTGKTITLEV<br>ESSDTIDNVKSKIQDKE<br>GIPPDQQRLIFAGKQLE<br>DGRTLSDYNIQKESTL<br>HLVLRLRGGYLFVQEL<br>ASKGEELFTGVVPILVE<br>LDGDVNGHKFSVRGE<br>GEGDATNGKLTLKFIC<br>TTGKLPVPWPTLVTTL<br>TYGVQCFSRYPDHMK<br>QHDFFKSAMPEGYVQ<br>ERTISFKDDGTYKTRA<br>EVKFEGDTLVNRIELK<br>GIDFKEDGNILGHKLE<br>YNFNSHNVYITADKQK<br>NGIKANFKIRHNVEDG<br>SVQLADHYQQNTPIGD<br>GPVLLPDNHYLSTQSV<br>LSKDPNEKRDHMVLLE<br>FVTAAGITHGMDELYK<br>LEHHHHHHH |
| pACYC-CvTA-AlaDH | CvTA:<br>ATGGGCAGCAGCCATCACCATCATCACCACAGCCAGGATCCGAATT<br>CGATGCAGAAGCAGCGTACAACATCGCAATGGCGCGAACTTGACG<br>CCGCTCATCACCTGCATCCCTTCACCGATACCGCCTCCCTTAACCAG<br>GCCGGCGCGCGCGTGATGACACGTGGAGAAGGGGTGTATTTGTGG<br>GACTCGGAGGGAAATAAAATCATCGACGGTATGGCTGGATTATGG<br>TGTGTGAACGTTGGCTACGGTCGTAAGGACTTTGCCGAAGCGGCCC<br>GTCGTCAGATGGAAGAATTACCGTTCTACAATACTTTTTTCAAAAC<br>AACCCATCCTGCGGTCGTAGAGTTATCTTCATTATTGGCGGAAGTC<br>ACTCCAGCAGGGTTTGACCGCGTGTTTTATACAAATAGTGGATCAG<br>AATCGGTTGACACAATGATCCGTATGGTCCGTCGTTACTGGGACGT<br>CCAAGGCAAACCGGAGAAGAAGACGTTAATCGGCCGCTGGAATGG<br>TTATCACGGTTCGACCATTGGAGGTGCATCTCTTGGGGGCATGAAG<br>TATATGCATGAGCAGGGTGATTTGCCTATCCCTGGCATGGCGCACA<br>TCGAACAACCGTGGTGGTATAAGCACGGTAAAGACATGACGCCGG<br>ACGAGTTTGGAGTTGTCGCTGCGCGTTGGTTGGAAGAGAAGATCCT<br>GGAAATTGGGGCGGACAAGGTAGCCGCCTTCGTAGGAGAACCAAT<br>CCAAGGTGCCGGGGGAGTGATCGTCCCGCCAGCTACCTATTGGCCC<br>GAGATCGAGCGCATTTGCCGTAAATATGACGTATTGCTGGTTGCAG<br>ATGAGGTAATTTGTGGCTTCGGGCGCACCGGGGAGTGGTTCGGGCA<br>CCAACATTTCGGTTTTCAGCCGGACTTATTTACGGCGGCGAAGGGT<br>TTAAGCTCAGGTTATTTACCGATTGGGGCTGTGTTTGTGGGCAAGC<br>GTGTTGCCGAAGGCTTAATCGCGGGAGGCGACTTTAATCACGGATT<br>CACATACTCTGGACACCCGGTTTGTGCCGCAGTAGCTCACGCGAAT<br>GTAGCCGCATTACGTGACGAGGGAATCGTCCAGCGTGTGAAGGAC<br>GATATCGGCCCTTATATGCAGAAGCGCTGGCGCGAGACTTTTTCAC<br>GTTTTGAGCACGTAGACGATGTGCGTGGCGTAGGCATGGTACAGGC<br>CTTTACCTTAGTCAAAAATAAAGCTAAGCGCGAGTTGTTCCCAGAC<br>TTTGGCGAAATCGGAACGTTGTGTCGCGATATCTTTTTCGCAATAA<br>TCTTATCATGCGCGCTTGCGGGGATCATATTGTAAGTGCCCCGCCA<br>TTGGTGATGACTCGTGCCGAGGTAGATGAGATGTTAGCAGTCGCAG<br>AGCGCTGCCTTGAGGAGTTTGAGCAAACATTAAAAGCTCGCGGACT<br>TGCCTGA<br>AlaDH: | CvTA:<br><br>MGSSHHHHHHSQDPN<br>SMQKQRTTSQWRELD<br>AAHHLHPFTDTASLNQ<br>AGARVMTRGEGVYLW<br>DSEGNKIIDGMAGLWC<br>VNVGYGRKDFAEAAR<br>RQMEELPFYNTFFKTT<br>HPAVVELSSLLAEVTP<br>AGFDRVFYTNSGSESV<br>DTMIRMVRRYWDVQG<br>KPEKKTLIGRWNGYH<br>GSTIGGASLGGMKYM<br>HEQGDLPIPGMAHIEQ<br>PWWYKHGKDMTPDEF<br>GVVAARWLEEKILEIG<br>ADKVAAFVGEPIQGAG<br>GVIVPPATYWPEIERIC<br>RKYDVLLVADEVICGF<br>GRTGEWFGHQHFGFQ<br>PDLFTAAKGLSSGYLPI<br>GAVFVGKRVAEGLIAG<br>GDFNHGFTYSGHPVCA<br>AVAHANVAALRDEGI<br>VQRVKDDIGPYMQKR<br>WRETFSRFEHVDDVRG<br>VGMVQAFTLVKNKAK<br>RELFPDFGEIGTLCRDI<br>FFRNNLIMRACGDHIV<br>SAPPLVMTRAEVDEML<br>AVAERCLEEFEQTLKA<br>RGLA* |

| | | |
|---|---|---|
| | ATGGGCAGCAGCCATCACCATCATCACCACATGATCATAGGGGTTC<br>CTAAAGAGATAAAAAACAATGAAAACCGTGTCGCATTAACACCCG<br>GGGGCGTTTCTCAGCTCATTTCAAACGGCCACCGGGTGCTGGTTGA<br>AACAGGCGCGGGCCTTGGAAGCGGATTTGAAAATGAAGCCTATGA<br>GTCAGCAGGAGCGGAAATCATTGCTGATCCGAAGCAGGTCTGGGA<br>CGCCGAAATGGTCATGAAAGTAAAAGAACCGCTGCCGGAAGAATA<br>TGTTTATTTTCGCAAAGGACTTGTGCTGTTTACGTACCTTCATTTAG<br>CAGCTGAGCCTGAGCTTGCACAGGCCTTGAAGGATAAAGGAGTAA<br>CTGCCATCGCATATGAAACGGTCAGTGAAGGCCGGACATTGCCTCT<br>TCTGACGCCAATGTCAGAGGTTGCGGGCAGAATGGCAGCGCAAAT<br>CGGCGCTCAATTCTTAGAAAAGCCTAAAGGCGGAAAAGGCATTCT<br>GCTTGCCGGGGTGCCTGGCGTTTCCCGCGGAAAAGTAACAATTATC<br>GGAGGAGGCGTTGTCGGGACAAACGCGGCGAAAATGGCTGTCGGC<br>CTCGGTGCAGATGTGACGATCATTGACTTAAACGCAGACCGCTTGC<br>GCCAGCTTGATGACATCTTCGGCCATCAGATTAAAACGTTAATTTC<br>TAATCCGGTCAATATTGCTGATGCTGTGGCGGAAGCGGATCTCCTC<br>ATTTGCGCGGTATTAATTCCGGGTGCTAAAGCTCCGACTCTTGTCAC<br>TGAGGAAATGGTAAAACAAATGAAACCCGGTTCAGTTATTGTTGAT<br>GTAGCGATCGACCAAGGCGGCATCGTCGAAACTGTCGACCATATCA<br>CAACACATGATCAGCCAACATATGAAAAACACGGGGTTGTGCATT<br>ATGCTGTAGCGAACATGCCAGGCGCAGTCCCTCGTACATCAACAAT<br>CGCCCTGACTAACGTTACTGTTCCATACGCGCTGCAAATCGCGAAC<br>AAAGGGGCAGTAAAAGCGCTCGCAGACAATACGGCACTGAGAGCG<br>GGTTTAAACACCGCAAACGGACACGTGACCTATGAAGCTGTAGCA<br>AGAGATCTAGGCTATGAGTATGTTCCTGCCGAGAAAGCTTTACAGG<br>ATGAATCATCTGTGGCGGGTGCTTAA | AlaDH:<br>MGSSHHHHHHMIIGVP<br>KEIKNNENRVALTPGG<br>VSQLISNGHRVLVETG<br>AGLGSGFENEAYESAG<br>AEIIADPKQVWDAEMV<br>MKVKEPLPEEYVYFRK<br>GLVLFTYLHLAAEPEL<br>AQALKDKGVTAIAYET<br>VSEGRTLPLLTPMSEV<br>AGRMAAQIGAQFLEKP<br>KGGKGILLAGVPGVSR<br>GKVTIIGGGVVGTNAA<br>KMAVGLGADVTIIDLN<br>ADRLRQLDDIFGHQIK<br>TLISNPVNIADAVAEA<br>DLLICAVLIPGAKAPTL<br>VTEEMVKQMKPGSVI<br>VDVAIDQGGIVETVDH<br>ITTHDQPTYEKHGVVH<br>YAVANMPGAVPRTSTI<br>ALTNVTVPYALQIANK<br>GAVKALADNTALRAG<br>LNTANGHVTYEAVAR<br>DLGYEYVPAEKALQD<br>ESSVAGA* |
| pZE-MaCAR-sfp | MaCAR:<br>atgggcagcagccatcaccatcatcaccacATGAGCACGGCCACCCATGATGAACGC<br>TTAGACCGCCGCGTGCACGAGCTGATTGCAACTGACCCTCAGTTTG<br>CGGCGGCTCAACCTGATCCCGCAATTACGGCTGCTCTTGAGCAACC<br>TGGTTTGCGTTTACCACAGATCATTCGTACAGTCCTGGACGGATAT<br>GCCGACCGTCCGGCATTGGGACAGCGCGTGGTCGAGTTCGTTACCG<br>ACGCTAAAACCGGTCGTACTAGCGCTCAACTCCTGCCGCGCTTTGA<br>GACCATCACATACGGAGAGGTGGCACAACGTGTTTCAGCCTTAGGT<br>CGTGCACTGAGCGACGATGCGGTTCACCCGGGCGATCGTGTTTGTG<br>TGTTAGGGTTCAATAGCGTTGATTATGCAACTATCGACATGGCGCT<br>CGGTGCAATCGGGGCGGTGAGCGTTCCACTCCAAACGTCAGCTGCC<br>ATTTCATCATTGCAGCCCATCGTAGCTGAGACTGAGCCAACACTTA<br>TCGCCAGCTCTGTGAATCAACTGTCCGATGCGGTCCAACTGATCAC<br>TGGCGCTGAGCAAGCCCCGACGCGCCTCGTCGTGTTCGACTACCAC<br>CCACAAGTGGACGACCAACGTGAAGCAGTTCAAGACGCAGCTGCC<br>CGTTTGTCAGGTACAGGTGTCGCGGTGCAAACATTGGCGGAGCTTT<br>TAGAACGCGGTAAGGACCTTCCAGCCGTTGCTGAGCCACCCGCAGA<br>TGAGGATTCTTTGGCGTTATTGATCTACACTTCCGGTTCGACGGGTG<br>CCCCCAAAGGCGCGATGTACCCTCAGTCTAACGTGGGGAAGATGTG<br>GCGTCGTGGTTCAAAGAACTGGTTTGGGGAGTCCGCAGCGTCTATT<br>ACCTTAAACTTTATGCCCATGTCCCACGTGATGGGGCGCTCCATTCT<br>GTACGGAACTTTAGGAAATGGCGGGACCGCCTACTTTGCAGCCCGT<br>TCGGATTTAAGCACCCTGTTAGAAGACCTGGAATTAGTACGCCCTA<br>CGGAGCTTAATTTTGTACCTCGTATCTGGGAGACTTTATATGGCGA<br>GTTCCAACGCCAAGTGGAGCGTCGCCTTTCAGAGGCCGGGGACGCC<br>GGCGAGCGTCGCGCGGTTGAAGCCGAAGTATTGGCCGAACAACGT<br>CAATACTTGTTGGGCGGCCGCTTCACGTTCGCTATGACGGGATCGG<br>CGCCGATTAGCCCCGAGCTGCGTAACTGGGTAGAGAGTCTGTTGGA<br>GATGCACTTGATGGATGGTTACGGCAGTACTGAGGCAGGTATGGTG<br>CTCTTTGACGGAGAAATTCAGCGTCCGCCTGTCGTGGACTACAAGC<br>TGGTAGATGTACCTGACTTAGGCTACTTTTCCACTGACCGTCCGCAC<br>CCGCGCGGTGAATTACTGTTACGTACTGAGAACATGTTTCCAGGCT<br>ATTATAAGCGTGCAGAAACGACTGCTGGGGTTTTCGACGAGGACG<br>GGTATTACCGCACTGGCGACGTGTTCGCCGAGATCGCGCCCGACCG<br>CTTGGTGTATGTCGATCGCCGCAATAATGTGTTGAAGCTGGCGCAG | MaCAR:<br><br>MGSSHHHHHHMSTAT<br>HDERLDRRVHELIATD<br>PQFAAAQPDPAITAAL<br>EQPGLRLPQIIRTVLDG<br>YADRPALGQRVVEFVT<br>DAKTGRTSAQLLPRFE<br>TITYGEVAQRVSALGR<br>ALSDDAVHPGDRVCV<br>LGFNSVDYATIDMALG<br>AIGAVSVPLQTSAAISS<br>LQPIVAETEPTLIASSV<br>NQLSDAVQLITGAEQA<br>PTRLVVFDYHPQVDDQ<br>REAVQDAAARLSGTG<br>VAVQTLAELLERGKDL<br>PAVAEPPADEDSLALLI<br>YTSGSTGAPKGAMYP<br>QSNVGKMWRRGSKN<br>WFGESAASITLNFMPM<br>SHVMGRSILYGTLGNG<br>GTAYFAARSDLSTLLE<br>DLELVRPTELNFVPRI<br>WETLYGEFQRQVERRL<br>SEAGDAGERRAVEAE<br>VLAEQRQYLLGGRFTF<br>AMTGSAPISPELRNWV<br>ESLLEMHLMDGYGSTE<br>AGMVLFDGEIQRPPVV<br>DYKLVDVPDLGYFSTD<br>RPHPRGELLLRTENMF<br>PGYYKRAETTAGVFDE<br>DGYYRTGDVFAEIAPD<br>RLVYVDRRNNVLKLA |

| | | |
|---|---|---|
| | GGTGAGTTCGTGACGTTAGCCAAGCTCGAGGCGGTATTCGGGAACA<br>GCCCTCTGATTCGTCAGATCTACGTCTACGGCAATTCGGCCCAGCC<br>ATATCTTCTTGCAGTTGTTGTTCCCACAGAAGAGGCACTGGCGAGT<br>GGGGACCCAGAGACCTTAAAGCCAAAGATCGCCGACTCACTCCAG<br>CAAGTAGCGAAAGAGGCCGGATTGCAATCCTATGAGGTGCCCCGT<br>GACTTTATTATCGAAACGACACCGTTTTCCTTAGAGAACGGGCTGC<br>TTACAGGCATCCGTAAGTTGGCCTGGCCAAAACTCAAGCAACATTA<br>TGGTGAGCGCTTAGAGCAAATGTATGCAGATCTGGCGGCGGGGCA<br>AGCAGATGAGTTAGCAGAACTCCGTCGCAATGGCGCCCAAGCGCC<br>TGTTCTGCAAACTGTGTCCCGCGCAGCAGGCGCAATGTTGGGGTCA<br>GCGGCATCTGACCTGAGTCCCGACGCACATTTTACGGATTTAGGCG<br>GCGATTCGCTCAGCGCACTCACGTTTGGCAATTTGCTGCGTGAGAT<br>CTTTGATGTCGACGTGCCAGTAGGTGTAATTGTCAGTCCAGCGAAT<br>GATCTTGCAGCCATCGCGTCGTATATTGAGGCTGAACGCCAGGGTT<br>CGAAACGCCCTACCTTCGCGAGCGTTCATGGGCGTGACGCAACCGT<br>TGTGCGTGCGGCTGACCTGACTCTGGACAAGTTTCTTGATGCTGAT<br>ACCCTTGCAAGTGCGCCCAATTTACCAAAACCAGCTACCGAAGTGC<br>GCACAGTCCTGTTAACCGGCGCGACAGGCTTCTTAGGTCGCTATCT<br>GGCCCTTGAGTGGCTGGAACGCATGGACATGGTCGATGGAAAGGT<br>CATTGCCTTAGTTCGCGCACGCAGCGACGAGGAAGCCCGTGCGCGT<br>CTCGACAAGACGTTTGACTCGGGTGACCCGAAGCTTCTTGCCCACT<br>ACCAACAATTAGCCGCCGACCACTTGGAAGTCATTGCAGGTGATAA<br>GGGCGAGGCGAACCTTGGGCTGCGTCAAGATGTATGGCAGCGTCT<br>GGCCGATACAGTTGATGTCATCGTTGACCCGGCAGCATTAGTGAAT<br>CACGTGCTTCCTTATAGTGAGTTATTTGGACCGAATGCCTTAGGCA<br>CGGCGGAATTAATTCGTTTAGCACTTACTAGCAAGCAGAAACCGTA<br>TACTTACGTCAGTACCATCGGCGTCGGTGATCAAATTGAGCCTGGA<br>AAATTCGTCGAGAATGCTGATATCCGTCAAATGTCCGCAACACGTG<br>CGATCAACGATTCTTATGCGAACGGTTACGGTAACAGTAAATGGGC<br>TGGAGAGGTCCTCTTGCGTGAGGCTCATGATCTGTGCGGTCTGCCC<br>GTCGCGGTGTTTCGTTGTGATATGATTCTGGCCGACACTACATATGC<br>CGGTCAATTAAATCTGCCAGATATGTTCACGCGCCTTATGCTGTCTT<br>TAGTTGCCACGGGAATCGCACCAGGTAGCTTTTACGAACTGGACGC<br>AGATGGCAATCGCCAACGTGCGCACTACGATGGGTTGCCGGTCGA<br>GTTCATCGCAGCGGCCATTAGTACCCTGGGTTCGCAGATCACAGAC<br>AGCGATACCGGCTTCCAAACATACCACGTAATGAATCCTTACGACG<br>ACGGGATTGGTCTTGACGAATATGTCGACTGGTTAGTGGACGCGGG<br>ATACAGCATTGAGCGCATTGCAGATTATTCTGAATGGCTTCGTCGT<br>TTTGAAACCAGTTTACGTGCACTGCCGGATCGCCAACGCCAGTATT<br>CACTTCTTCCCTTACTGCACAACTACCGTACGCCTGAGAAGCCTATT<br>AACGGCAGCATTGCACCTACCGACGTTTTCCGCGCTGCCGTCCAGG<br>AAGCGAAAATCGGTCCGGATAAGGACATCCCACATGTGAGCCCGC<br>CGGTCATTGTCAAGTACATTACGGATTTGCAACTGCTGGGGTTGCT<br>CTAA<br>sfp:<br>atgaaaatctatggcatttacatggatcgtccgctgagtcaggaagaaaacgaacgctttatgaccttcatcagcc<br>cggaaaaacgtgaaaaatgccgtcgcttttatcataaagaagatgcacaccgcacgctgctgggcgatgtgctg<br>gttcgtagcgtgatctctcgccagtatcagctggataaatctgatattcgtttcagtacccaggaatacggtaaacc<br>gtgtattccggatctgccggatgcacattttaatatcagccactctggccgctgggttattggtgcgttcgattctca<br>gccgattggtatcgatattgaaaaaacgaaaccgatcagtctggaaattgccaaacgtttctttagcaaaaccgaa<br>tattctgatctgctggcaaaagataaagatgaacagacggattacttttaccatctgtggagtatgaaagaatctttt<br>atcaaacaggaaggcaaaggtctgagcctgccgctggatagttttagcgtgcgcctgcatcaggatggccagg<br>tttctatcgaactgccggattctcacagtccgtgctatattaaaacctacgaagttgatccgggctataaaatggcc<br>gtttgtgcggcccacccggatttcccggaagatattacgatggtgagctacgaagaactgctgtaa | QGEFVTLAKLEAVFGN<br>SPLIRQIYVYGNSAQPY<br>LLAVVVPTEEALASGD<br>PETLKPKIADSLQQVA<br>KEAGLQSYEVPRDFIIE<br>TTPFSLENGLLTGIRKL<br>AWPKLKQHYGERLEQ<br>MYADLAAGQADELAE<br>LRRNGAQAPVLQTVSR<br>AAGAMLGSAASDLSP<br>DAHFTDLGGDSLSALT<br>FGNLLREIFDVDVPVG<br>VIVSPANDLAAIASYIE<br>AERQGSKRPTFASVHG<br>RDATVVRAADLTLDK<br>FLDADTLASAPNLPKP<br>ATEVRTVLLTGATGFL<br>GRYLALEWLERMDMV<br>DGKVIALVRARSDEEA<br>RARLDKTFDSGDPKLL<br>AHYQQLAADHLEVIA<br>GDKGEANLGLRQDVW<br>QRLADTVDVIVDPAAL<br>VNHVLPYSELFGPNAL<br>GTAELIRLALTSKQKP<br>YTYVSTIGVGDQIEPG<br>KFVENADIRQMSATRA<br>INDSYANGYGNSKWA<br>GEVLLREAHDLCGLPV<br>AVFRCDMILADTTYAG<br>QLNLPDMFTRLMLSLV<br>ATGIAPGSFYELDADG<br>NRQRAHYDGLPVEFIA<br>AAISTLGSQITDSDTGF<br>QTYHVMNPYDDGIGL<br>DEYVDWLVDAGYSIE<br>RIADYSEWLRRFETSL<br>RALPDRQRQYSLLPLL<br>HNYRTPEKPINGSIAPT<br>DVFRAAVQEAKIGPDK<br>DIPHVSPPVIVKYITDL<br>QLLGLL*<br><br>sfp:<br>MKIYGIYMDRPLSQEE<br>NERFMTFISPEKREKCR<br>RFYHKEDAHRTLLGD<br>VLVRSVISRQYQLDKS<br>DIRFSTQEYGKPCIPDL<br>PDAHFNISHSGRWVIG<br>AFDSQPIGIDIEKTKPIS<br>LEIAKRFFSKTEYSDLL<br>AKDKDEQTDYFYHLW<br>SMKESFIKQEGKGLSLP |

| | | |
|---|---|---|
| | | LDSFSVRLHQDGQVSI<br>ELPDSHSPCYIKTYEVD<br>PGYKMAVCAAHPDFP<br>EDITMVSYEELL* |
| pET-28a(+)-<br>Turbo PETase | atgAGCAATCCGTATCAGCGTGGTCCGAATCCGACACGTAGCGCACT<br>GACCACCGATGGTCCGTTTAGCGTTGCAACCTATAGCGTTAGCCGT<br>CTGAGCGTTAGCGGTTTTGGTGGTGGTGTTATCTATTATCCGACCGG<br>TACAACCCTGACCTTTGGTGGTATTGCAATGAGTCCGGGTTATACC<br>GCAGATGCAAGCAGCCTGGCACTGCTGGGTCGTCGTCTGGCAAGCC<br>ATGGTTTTGTTGTTATTGTGATTAATACCAACAGCCGTCTGGATTTT<br>CCGGATAGCCGTGCAAGCCAGCTGAGCGCAGCACTGAATTATCTGC<br>GTACCAGCAGTCCGAGCGCAGTTCGTGCACGTCTGGATGCAAATCG<br>TCTGGCCGTTGCAGGTCATAGCATGGGTGGTGGCGCAACCCTGCGT<br>ATTAGCGAGCAGATTCCGACACTGAAAGCCGGTGTTCCGCTGACAC<br>CGTGGCATACCGATAAAACCTTTAATACACCGGTTCCGCAGCTGAT<br>TGTTGGTGCAGAACGTGATACCGTTGCACCGGTTAGCCAGAGCGCA<br>ATTCCGATTTATCAGAATCTGCCGAGCACCACACCGAAAGTTTATG<br>TTGAACTGAAGAATGCGACCCATACCGCACCGAATAGCCCGAATG<br>CATGCATTAGCGTTTATACCATTAGCTGGATGAAACTGTGGGTTGA<br>TAATGATACCCGTTATCGTCAGTTTCTGTGCAATGTTAATGATCCGT<br>GCCTGAGCGATTTTCGTAGCAATAATCGTCATTGTCAGCTCGAGcacc<br>accaccaccaccactga | MSNPYQRGPNPTRSAL<br>TTDGPFSVATYSVSRLS<br>VSGFGGGVIYYPTGTT<br>LTFGGIAMSPGYTADA<br>SSLALLGRRLASHGFV<br>VIVINTNSRLDFPDSRA<br>SQLSAALNYLRTSSPSA<br>VRARLDANRLAVAGH<br>SMGGGATLRISEQIPTL<br>KAGVPLTPWHTDKTF<br>NTPVPQLIVGAERDTV<br>APVSQSAIPIYQNLPST<br>TPKVYVELKNATHTAP<br>NSPNACISVYTISWMK<br>LWVDNDTRYRQFLCN<br>VNDPCLSDFRSNNRHC<br>QLEHHHHHH* |
| pET-28a(+)-<br>Bhr PETase | atgAGCAATCCGTATCAGCGTGGTCCGAATCCGACACGTAGCGCACT<br>GACCACCGATGGTCCGTTTAGCGTTGCAACCTATAGCGTTAGCCGT<br>CTGAGCGTTAGCGGTTTTGGTGGTGGTGTTATCTATTATCCGACCGG<br>TACAACCCTGACCTTTGGTGGTATTGCAATGAGTCCGGGTTATACC<br>GCAGATGCAAGCAGCCTGGCATGGCTGGGTCGTCGTCTGGCAAGCC<br>ATGGTTTTGTTGTTATTGTGATTAATACCAACAGCCGTCTGGATTTT<br>CCGGATAGCCGTGCAAGCCAGCTGAGCGCAGCACTGAATTATCTGC<br>GTACCAGCAGTCCGAGCGCAGTTCGTGCACGTCTGGATGCAAATCG<br>TCTGGCCGTTGCAGGTCATAGCATGGGTGGTGGCGCAACCCTGCGT<br>ATTAGCGAGCAGATTCCGACACTGAAAGCCGGTGTTCCGCTGACAC<br>CGTGGCATACCGATAAAACCTTTAATACACCGGTTCCGCAGCTGAT<br>TGTTGGTGCAGAAGCAGATACCGTTGCACCGGTTAGCCAGCATGCA<br>ATTCCGTTTTATCAGAATCTGCCGAGCACCACACCGAAAGTTTATG<br>TTGAACTGGATAATGCGACCCATTTTGCACCGAATAGCCCGAATGC<br>AGCAATTAGCGTTTATACCATTAGCTGGATGAAACTGTGGGTTGAT<br>AATGATACCCGTTATCGTCAGTTTCTGTGCAATGTTAATGATCCGGC<br>ACTGAGCGATTTTCGTAGCAATAATCGTCATTGTCAGCTCGAGcacca<br>ccaccaccaccactga | MSNPYQRGPNPTRSAL<br>TTDGPFSVATYSVSRLS<br>VSGFGGGVIYYPTGTT<br>LTFGGIAMSPGYTADA<br>SSLAWLGRRLASHGFV<br>VIVINTNSRLDFPDSRA<br>SQLSAALNYLRTSSPSA<br>VRARLDANRLAVAGH<br>SMGGGATLRISEQIPTL<br>KAGVPLTPWHTDKTF<br>NTPVPQLIVGAEADTV<br>APVSQHAIPFYQNLPST<br>TPKVYVELDNATHFAP<br>NSPNAAISVYTISWMK<br>LWVDNDTRYRQFLCN<br>VNDPALSDFRSNNRHC<br>QLEHHHHHH* |
| pET-28a(+)-<br>TS PETase | atgCAGACCAATCCATACGCTCGTGGTCCAAATCCGACCGCCGCAAG<br>CCTGGAAGCAAGCGCAGGTCCATTTACCGTTCGCAGCTTTACCGTT<br>AGCCGTCCAAGCGGTTATGGTGCAGGTACCGTTTATTATCCGACCA<br>ATGCAGGTGGCACCGTTGGTGCAATTGCTATTGTTCCGGGTTATAC<br>CGCCCGCCAGAGCAGCATTAAATGGTGGGGTCCGCGCCTGGCCAGT<br>CATGGTTTTGTTGTTATTACCATTGATACCAATAGCACCCTGGATCA<br>GCCGGAAAGCCGTTCAAGTCAGCAGATGGCAGCACTGCGTCAGGT<br>GGCGTCTCTGAATGGTACTAGTAGCAGTCCGATTTATGGTAAAGTT<br>GATACCGCACGTATGGGCGTTATGGGTTGGAGTATGGGTGGTGGTG<br>GTAGTCTGATTAGTGCCGCTAATAATCCGAGCCTGAAAGCAGCGGC<br>ACCGCAGGCACCGTGGCATAGCAGTACCAACTTTAGTAGCGTTACG<br>GTTCCGACCCTGATTTTTGCTTGTGAAAATGATAGCATTGCACCGGT<br>TAATAGCAGCGCACTGCCGATTTATGATTCAATGAGCcgcAATGCAA<br>AACAGTTTCTGGAAATTtgtGGCGGTAGCCATTCTTGTGCCAATAGTG<br>GTAATAGCAATCAGGCACTGATTGGTAAAAAGGGTGTTGCCTGGAT<br>GAAACGTTTTATGGATAACGATACCCGTTATAGCACCTTTGCATGT<br>GAAAATCCGAATAGTACCGCCGTTTGTGATTTTCGCACCGCAAATT<br>GCAGTCTCGAGCACCACCACCACCACCACTGA | MQTNPYARGPNPTAAS<br>LEASAGPFTVRSFTVSR<br>PSGYGAGTVYYPTNA<br>GGTVGAIAIVPGYTAR<br>QSSIKWWGPRLASHGF<br>VVITIDTNSTLDQPESR<br>SSQQMAALRQVASLN<br>GTSSSPIYGKVDTARM<br>GVMGWSMGGGGSLIS<br>AANNPSLKAAAPQAP<br>WHSSTNFSSVTVPTLIF<br>ACENDSIAPVNSSALPI<br>YDSMSRNAKQFLEICG<br>GSHSCANSGNSNQALI<br>GKKGVAWMKRFMDN<br>DTRYSTFACENPNSTA<br>VCDFRTANCSLEHHHH<br>HH* |

| | | |
|---|---|---|
| pET-28a(+)-<br>Thermo<br>PETase | atgCAGACCAATCCATACGCTCGTGGTCCAAATCCGACCGCCGCAAG<br>CCTGGAAGCAAGCGCAGGTCCATTTACCGTTCGCAGCTTTACCGTT<br>AGCCGTCCAAGCGGTTATGGTGCAGGTACCGTTTATTATCCGACCA<br>ATGCAGGTGGCACCGTTGGTGCAATTGCTATTGTTCCGGGTTATAC<br>CGCCCGCCAGAGCAGCATTAAATGGTGGGGTCCGCGCCTGGCCAGT<br>CATGGTTTTGTTGTTATTACCATTGATACCAATAGCACCCTGGATCA<br>GCCGGAAAGCCGTTCAAGTCAGCAGATGGCAGCACTGCGTCAGGT<br>GGCGTCTCTGAATGGTACTAGTAGCAGTCCGATTTATGGTAAAGTT<br>GATACCGCACGTATGGGCGTTATGGGTTGGAGTATGGGTGGTGGTG<br>GTAGTCTGATTAGTGCCGCTAATAATCCGAGCCTGAAAGCAGCGGC<br>ACCGCAGGCACCGTGGCATAGCAGTACCAACTTTAGTAGCGTTACG<br>GTTCCGACCCTGATTTTTGCTTGTGAAAATGATAGCATTGCACCGGT<br>TAATAGCAGCGCACTGCCGATTTATGATTCAATGAGCcgcAATGCAA<br>AACAGTTTCTGGAAATTaacGGCGGTAGCCATTCTTGTGCCAATAGT<br>GGTAATAGCAATCAGGCACTGATTGGTAAAAAGGGTGTTGCCTGG<br>ATGAAACGTTTTATGGATAACGATACCCGTTATAGCACCTTTGCAT<br>GTGAAAATCCGAATAGTACCGCCGTTAGTGATTTTCGCACCGCAAA<br>TTGCAGTCTCGAGCACCACCACCACCACCACTGA | MQTNPYARGPNPTAAS<br>LEASAGPFTVRSFTVSR<br>PSGYGAGTVYYPTNA<br>GGTVGAIAIVPGYTAR<br>QSSIKWWGPRLASHGF<br>VVITIDTNSTLDQPESR<br>SSQQMAALRQVASLN<br>GTSSSPIYGKVDTARM<br>GVMGWSMGGGGSLIS<br>AANNPSLKAAAPQAP<br>WHSSTNFSSVTVPTLIF<br>ACENDSIAPVNSSALPI<br>YDSMSRNAKQFLEING<br>GSHSCANSGNSNQALI<br>GKKGVAWMKRFMDN<br>DTRYSTFACENPNSTA<br>VSDFRTANCSLEHHHH<br>HH* |
| pET-28a(+)-<br>Dura PETase | atgCAGACCAATCCATACGCTCGTGGTCCAAATCCGACCGCCGCAAG<br>CCTGGAAGCAAGCGCAGGTCCATTTACCGTTCGCAGCTTTACCGTT<br>AGCCGTCCAAGCGGTTATGGTGCAGGTACCGTTTATTATCCGACCA<br>ATGCAGGTGGCACCGTTGGTGCAATTGCTATTGTTCCGGGTTATAC<br>CGCCCGCCAGAGCAGCATTAAATGGTGGGGTCCGCGCCTGGCCAGT<br>CATGGTTTTGTTGTTATTACCATTGATACCAATAGCACCttcGATtatCC<br>GagcAGCCGTTCAAGTCAGCAGATGGCAGCACTGCGTCAGGTGGCG<br>TCTCTGAATGGTgatAGTAGCAGTCCGATTTATGGTAAAGTTGATAC<br>CGCACGTATGGGCGTTATGGGTcacAGTATGGGTGGTGGTgcgAGTCT<br>GcgtAGTGCCGCTAATAATCCGAGCCTGAAAGCAGCGataCCGCAGGC<br>ACCGTGGgacAGCcagACCAACTTTAGTAGCGTTACGGTTCCGACCCT<br>GATTTTTGCTTGTGAAAATGATAGCATTGCACCGGTTAATAGCcatGC<br>ACTGCCGATTTATGATTCAATGAGCcgcAATGCAAAACAGTTTCTGG<br>AAATTaacGGCGGTAGCCATTCTTGTGCCAATAGTGGTAATAGCAAT<br>CAGGCACTGATTGGTAAAAAGGGTGTTGCCTGGATGAAACGTTTTA<br>TGGATAACGATACCCGTTATAGCACCTTTGCATGTGAAAATCCGAA<br>TAGTACCGCCGTTAGTGATTTTCGCACCGCAAATTGCAGTCTCGAG<br>CACCACCACCACCACCACTGA | MQTNPYARGPNPTAAS<br>LEASAGPFTVRSFTVSR<br>PSGYGAGTVYYPTNA<br>GGTVGAIAIVPGYTAR<br>QSSIKWWGPRLASHGF<br>VVITIDTNSTFDYPSSR<br>SSQQMAALRQVASLN<br>GDSSSPIYGKVDTARM<br>GVMGHSMGGGASLRS<br>AANNPSLKAAIPQAPW<br>DSQTNFSSVTVPTLIFA<br>CENDSIAPVNSHALPIY<br>DSMSRNAKQFLEINGG<br>SHSCANSGNSNQALIG<br>KKGVAWMKRFMDND<br>TRYSTFACENPNSTAV<br>SDFRTANCSLEHHHHH<br>H* |
| pET-28a(+)-<br>FAST PETase<br>and<br>pACYC-<br>FAST PETase | atgCAGACCAATCCATACGCTCGTGGTCCAAATCCGACCGCCGCAAG<br>CCTGGAAGCAAGCGCAGGTCCATTTACCGTTCGCAGCTTTACCGTT<br>AGCCGTCCAAGCGGTTATGGTGCAGGTACCGTTTATTATCCGACCA<br>ATGCAGGTGGCACCGTTGGTGCAATTGCTATTGTTCCGGGTTATAC<br>CGCCCGCCAGAGCAGCATTAAATGGTGGGGTCCGCGCCTGGCCAGT<br>CATGGTTTTGTTGTTATTACCATTGATACCAATAGCACCCTGGATCA<br>GCCGGAAAGCCGTTCAAGTCAGCAGATGGCAGCACTGCGTCAGGT<br>GGCGTCTCTGAATGGTACTAGTAGCAGTCCGATTTATGGTAAAGTT<br>GATACCGCACGTATGGGCGTTATGGGTTGGAGTATGGGTGGTGGTG<br>GTAGTCTGATTAGTGCCGCTAATAATCCGAGCCTGAAAGCAGCGGC<br>ACCGCAGGCACCGTGGCATAGCAGTACCAACTTTAGTAGCGTTACG<br>GTTCCGACCCTGATTTTTGCTTGTGAAAATGATAGCATTGCACCGGT<br>TAATAGCAGCGCACTGCCGATTTATGATTCAATGAGCCAGAATGCA<br>AAACAGTTTCTGGAAATTAAAGGCGGTAGCCATTCTTGTGCCAATA<br>GTGGTAATAGCAATCAGGCACTGATTGGTAAAAAGGGTGTTGCCTG<br>GATGAAACGTTTTATGGATAACGATACCCGTTATAGCACCTTTGCA<br>TGTGAAAATCCGAATAGTACCGCCGTTAGTGATTTTCGCACCGCAA<br>ATTGCAGTCTCGAGCACCACCACCACCACCACTGA | MQTNPYARGPNPTAAS<br>LEASAGPFTVRSFTVSR<br>PSGYGAGTVYYPTNA<br>GGTVGAIAIVPGYTAR<br>QSSIKWWGPRLASHGF<br>VVITIDTNSTLDQPESR<br>SSQQMAALRQVASLN<br>GTSSSPIYGKVDTARM<br>GVMGWSMGGGGSLIS<br>AANNPSLKAAAPQAP<br>WHSSTNFSSVTVPTLIF<br>ACENDSIAPVNSSALPI<br>YDSMSQNAKQFLEIKG<br>GSHSCANSGNSNQALI<br>GKKGVAWMKRFMDN<br>DTRYSTFACENPNSTA<br>VSDFRTANCSLEHHHH<br>HH* |
| pET-28a(+)-<br>PES-H1<br>PETase | ATGGCGAACCCGTACGAGCGCGGGCCCGATCCCACCGAGTCGAGC<br>ATCGAGGCCGTCCGCGGGCCGTTCGCCGTGGCCCAGACGACGGTGT<br>CGAGGCTCCAGGCCGACGGCTTCGGCGGCGGGACCATCTACTACCC<br>GACCGACACGAGCCAGGGCACCTTCGGTGCGGTGGCGATCTCGCC<br>GGGGTTCACGGCGGGCCAGGAGAGCATCGCCTGGCTCGGCCCCCG | MANPYERGPDPTESSIE<br>AVRGPFAVAQTTVSRL<br>QADGFGGGTIYYPTDT<br>SQGTFGAVAISPGFTA |

| | | |
|---|---|---|
| | CATCGCGTCGCAGGGCTTCGTGGTGATCACGATCGACACGATCACG<br>CGCTTCGACTATCCCGACAGCCGGGGTCGCCAGCTGCAGGCCGCGC<br>TCGACCACCTGCGCACCAACAGCGTCGTGCGCAACCGGATCGACCC<br>GAACCGGATGGCGGTCATGGGCCACTCGATGGGCGGCGGCGGGGC<br>GCTGTCCGCCGCGGCGAACAACACGAGCCTCGAGGCCGCCATCCC<br>GCTGCAGGGCTGGCACACCCGGAAGAACTGGTCGAGCGTGCGGAC<br>GCCGACCCTGGTGGTCGGGGCCCAGCTCGACACCATCGCGCCGGTG<br>AGCTCGCACTCGGAGGCCTTCTACAACAGCCTGCCGAGCGACCTCG<br>ACAAGGCGTACATGGAGCTCCGCGGGGCCAGCCACCTCGTGTCGA<br>ACACGCCCGACACGACGACCGCGAAGTACAGCATCGCCTGGCTCA<br>AGCGGTTCGTCGACGACGACCTCCGCTACGAGCAGTTCCTGTGCCC<br>GGCGCCGGACGACTTCGCGATCTCCGAGTACCGCTCCACCTGCCCG<br>TTCCTCGAGCACCACCACCACCACCACTGA | GQESIAWLGPRIASQGF<br>VVITIDTITRFDYPDSR<br>GRQLQAALDHLRTNS<br>VVRNRIDPNRMAVMG<br>HSMGGGGALSAAANN<br>TSLEAAIPLQGWHTRK<br>NWSSVRTPTLVVGAQL<br>DTIAPVSSHSEAFYNSL<br>PSDLDKAYMELRGAS<br>HLVSNTPDTTTAKYSI<br>AWLKRFVDDDLRYEQ<br>FLCPAPDDFAISEYRST<br>CPFLEHHHHHH* |
| pET-28a(+)-<br>HOT PETase | ATGCAGACTAACCCCTATGCTCGCGGGCCGAATCCTACAGCGGCCT<br>CGTTGGAAGCCAGTGCAGGTCCCTTCACCGTACGTTCGTTTACTGTT<br>GCGCGTCCAGTGGGATATGGGGCTGGCACCGTCTATTATCCGACTA<br>ACGCCGGTGGTACTGTGGGCGCAATCGCCATTGTCCCCGGCTACAC<br>TGCAACTCAATCCTCGATTAATTGGTGGGGACCACGCTTGGCTAGC<br>CACGGGTTTGTTGTGATCACCATCGATACTAACAGTACGTTGGACA<br>AGCCAGAGAGTCGCAGCTCTCAGCAGATGGCGGCATTACGCCAGG<br>TGGCGAGCTTAAATGGGACGAGTTCAAGTCCAATTTATGGCAAGGT<br>CGATACCGCCCGTGGTGGAGTAATGGGCTGGAGTATGGGTGGCGG<br>CGGATCTCTTATCTCGGCAGCGAATAATCCAAGCCTGAAGGCAGCA<br>GCCGTTATGGCACCCTGGCATAGTAGTACGAACTTTTCGTCTGTTAC<br>GGTTCCTACGCTTATCTTTGCCTGTGAGAATGATAGGATCGCACCA<br>GTGAAGGAGTACGCCTTGCCAATTTACGACTCGATGTCGCTCAACG<br>CTAAACAGTTTCTTGAGATTTGTGGGGGCTCTCACTCTTGTGCCTGT<br>AGTGGGAATTCCAATCAAGCCCTTATCGGAATGAAGGGCGTTGCTT<br>GGATGAAGCGCTTCATGGACAATGACACTCGTTATTCACAGTTCGC<br>TTGCGAAAACCCAAACTCAACCGCCGTATGTGATTTTCGCACTGCT<br>AACTGCAGCCTCGAGCACCACCACCACCACCACTGA | MQTNPYARGPNPTAAS<br>LEASAGPFTVRSFTVA<br>RPVGYGAGTVYYPTN<br>AGGTVGAIAIVPGYTA<br>TQSSINWWGPRLASHG<br>FVVITIDTNSTLDKPES<br>RSSQQMAALRQVASL<br>NGTSSSPIYGKVDTAR<br>GGVMGWSMGGGGSLI<br>SAANNPSLKAAAVMA<br>PWHSSTNFSSVTVPTLI<br>FACENDRIAPVKEYAL<br>PIYDSMSLNAKQFLEIC<br>GGSHSCACSGNSNQAL<br>IGMKGVAWMKRFMD<br>NDTRYSQFACENPNST<br>AVCDFRTANCSLEHHH<br>HHH* |
| pET-28a(+)-<br>LCC-ICCG | atgAGCAACCCGTACCAGCGTGGCCCGAATCCGACCCGCAGCGCACT<br>GACCGCAGATGGCCCGTTTAGCGTGGCAACCTACACCGTCTCACGC<br>CTGTCAGTCTCGGGTTTTGGCGGTGGCGTGATTTATTACCCGACCG<br>GCACGTCTCTGACGTTCGGTGGCATCGCGATGAGTCCGGGTTATAC<br>CGCAGATGCTAGCTCTCTGGCATGGCTGGGTCGTCGCCTGGCTTCC<br>CATGGCTTTGTGGTTCTGGTGATTAACACGAATTCACGTTTCGATGG<br>CCCGGACAGCCGCGCCTCTCAGCTGAGTGCCGCCCTGAACTACCTG<br>CGTACCAGTTCCCCGAGCGCCGTTCGCGCACGTCTGGATGCAAATC<br>GTCTGGCGGTTGCCGGTCATTCTATGGGTGGCGGTGGCACCCTGCG<br>TATTGCAGAACAAAACCCGAGCCTGAAAGCGGCTGTCCCGCTGACC<br>CCGTGGCACACCGATAAAACGTTTAATACCAGTGTCCCGGTGCTGA<br>TTGTTGGCGCAGAAGCTGACACCGTGGCGCCGGTTTCGCAGCATGC<br>CATCCCGTTTTATCAAAACCTGCCGAGCACCACGCCGAAAGTTTAC<br>GTCGAACTGTGCAACGCATCGCACATTGCTCCGAATAGCAACAATG<br>CGGCCATTTCCGTTTATACGATCTCATGGATGAAACTGTGGGTCGA<br>TAATGACACCCGTTACCGCCAGTTCCTGTGTAATGTGAACGACCCG<br>GCTCTGTGCGACTTCCGCACCAATAATCGCCACTGCCAACTCGAGca<br>ccaccaccaccaccactga | MSNPYQRGPNPTRSAL<br>TADGPFSVATYTVSRL<br>SVSGFGGGVIYYPTGT<br>SLTFGGIAMSPGYTAD<br>ASSLAWLGRRLASHGF<br>VVLVINTNSRFDGPDS<br>RASQLSAALNYLRTSS<br>PSAVRARLDANRLAV<br>AGHSMGGGGTLRIAEQ<br>NPSLKAAVPLTPWHTD<br>KTFNTSVPVLIVGAEA<br>DTVAPVSQHAIPFYQN<br>LPSTTPKVYVELCNAS<br>HIAPNSNNAAISVYTIS<br>WMKLWVDNDTRYRQ<br>FLCNVNDPALCDFRTN<br>NRHCQLEHHHHHH* |
| pET-28a(+)-<br>LCC | atgagaaacagacaccaccaccatcaccatcgtgtgagacggcggagagagaatttgtatttccagagtaatcc<br>gtatcagagaggacccaacccgacgagaagtgccttgacagcggacggacccttcagcgtagccacctatac<br>ggtctctcgcttaagtgtgagtggattcggtggcggagtaatctactaccctacgggcacctctttgaccttcggt<br>ggcatcgctatgagcccgggctacaccgcggatgccagtagcctggcttggttgggacgtcggttggcgagtc<br>acgggtttgtggtcttggtcattaacacaaacagccggttcgactatcccgatagccgggctagtcaattatccgc<br>ggctctgaattacctccggaccagctctccgagtgctgttagagcacggctggatgcaaaccgtctcgcggtgg<br>ccggccatagcatgggtggaggcggtaccttgcggatcgcagaacagaacccgagcttaaaggccgcggttc<br>ctttgaccccgtggcatacggacaagacatttaataccagcgtcccggtcttaattgtgggatgtgaagcggata<br>ctgtggcgcccgtaagccaacatgcgattcctttctaccagaatctgccgagtactaccccgaaggtatatgtag<br>aactggataatgcctctcacttttgtcctaacagcaacaacgcagcaatcagtgtgtacacgattagttggatgaa | MRNRHHHHHHRVRRR<br>RENLYFQSNPYQRGPN<br>PTRSALTADGPFSVAT<br>YTVSRLSVSGFGGGVI<br>YYPTGTSLTFGGIAMSP<br>GYTADASSLAWLGRR<br>LASHGFVVLVINTNSR<br>FDYPDSRASQLSAALN<br>YLRTSSPSAVRARLDA<br>NRLAVAGHSMGGGGT<br>LRIAEQNPSLKAAVPLT |

| | | |
|---|---|---|
| | actgtgggtagacaacgatacacgataccgacagttcttgtgtaacgtgaacgacccggcgttgagcgatttcagaaccaataacagacattgtcagtaa | PWHTDKTFNTSVPVLIVGCEADTVAPVSQHAIPFYQNLPSTTPKVYVELDNASHFCPNSNNAAISVYTISWMKLWVDNDTRYRQFLCNVNDPALSDFRTNNRHCQ* |

**Table S4.**

Summary of PETases used in this study

| **PETase** | **Wild-type enzyme** | **Mutations** | **Optimum Temperature** |
|---|---|---|---|
| TS | IsPETase | R280A/S121E/D186H/N233C/S282C | 58 |
| Thermo | IsPETase | S121E/D186H/R280A | 40 |
| FAST | IsPETase | S121E/D186H/R224Q/N233K/R280A | 50 |
| PES | PESPETase | L92F/Q94Y | 70 |
| Dura | IsPETase | S214H−I168R−W159H−S188Q−R280A−A180I−G165A−Q119Y−L117F−T140D | 37 |
| BHR | BhrPETase | NA | |
| Turbo | BhrPETase | W104L/H218S/F222I/A209R/D238K/F243T/A251C/A281C | |
| LCC | LCC | A244C/A192C | 70 |
| ICCG | LCC | F243I/D238C/S283C/Y127G | |
| HOT | IsPETase | S121E/D186H/R280A/P181V/S207R/S214Y/Q119K/S213E/N233C/S282C/R90T/Q182M/N212K/R224L/S58A/S61V/K95N/M154G/N241C/K252M/T270Q | 60 |

**Supplemental Figures**

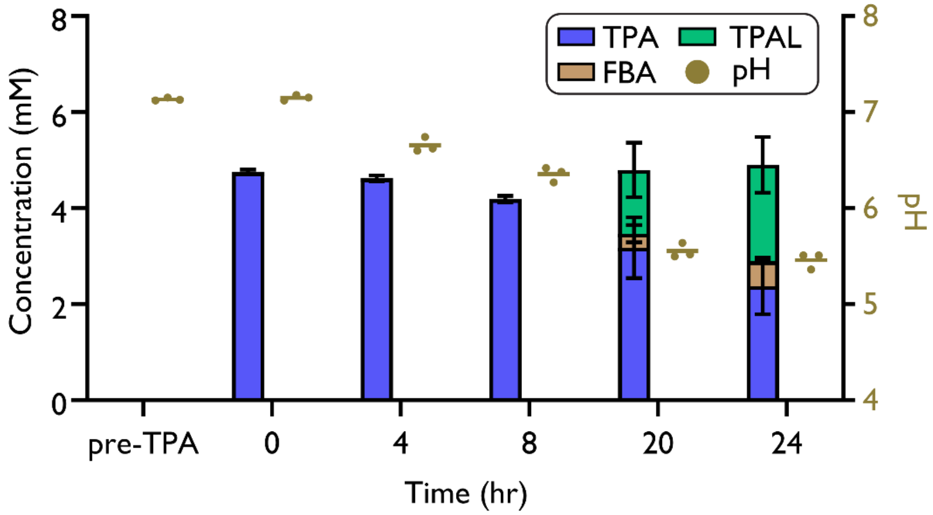


**Fig. S1. Evaluating pH effects on aldehyde production.**
RARE.Δ16 cells expressing MaCAR were cultured in MOPS media with 100 mM disodium phosphate at pH 7.4. Time course of analysis of the relationship between pH and TPAL production.

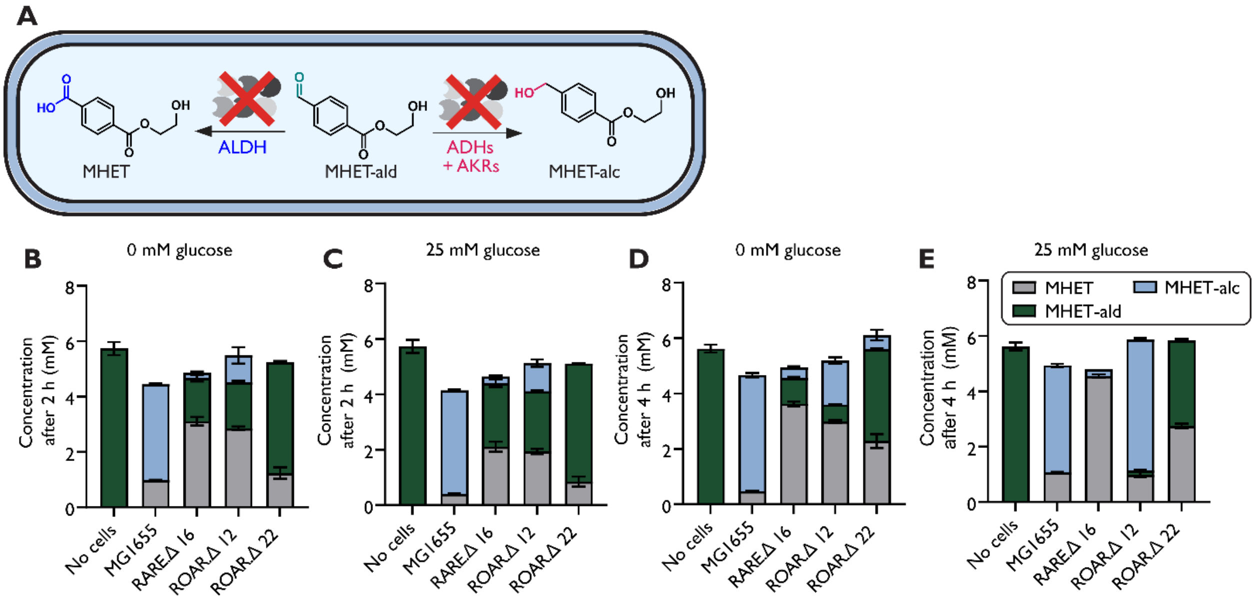


**Fig. S2. MHET-ald stability in resting cells.**
(**A**) *E. coli* resting cells of MG1655, RARE.Δ16, ROAR.Δ12 and ROAR.Δ22 were supplemented in 200 mM HEPES (**B**, **D**) without or (**C**, **E**) with 25 mM glucose at pH 7.5. MHET-ald stability was measured at (**B**, **C**) 2 h and (**D**, **E**) 4 h.

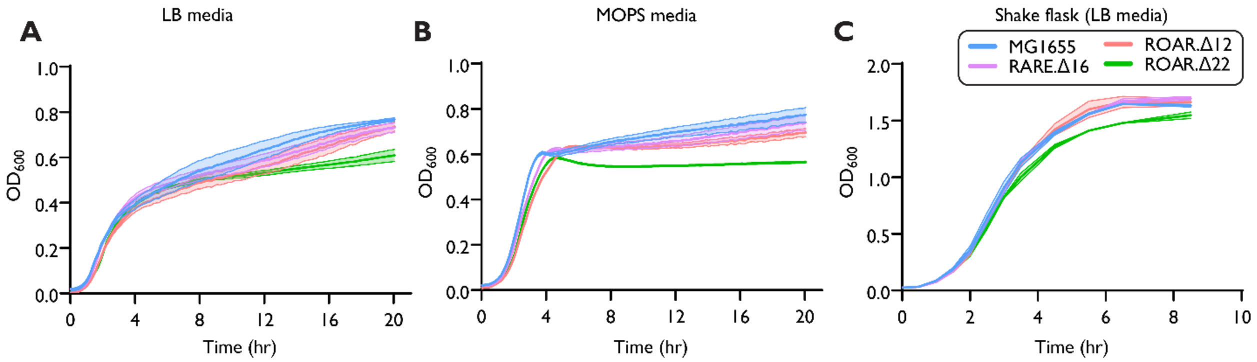


**Fig. S3**. **Fitness of engineered strains**.

Growth of wild-type *E. coli* MG1655, RARE.Δ16, ROAR.Δ12 and ROAR.Δ22 was monitored via $OD_{600}$ in 96-well plate for 12 h in (**A**) LB media and (**B**) MOPS media with 2 % glucose. (**C**) Growth of the previously mentioned strains in 250 mL shake flasks with 50 mL of LB media. Growth was tracked every 0.5 h in duplicate via spectrophotometer measurements to measure optical density at 600 nm ($OD_{600}$).

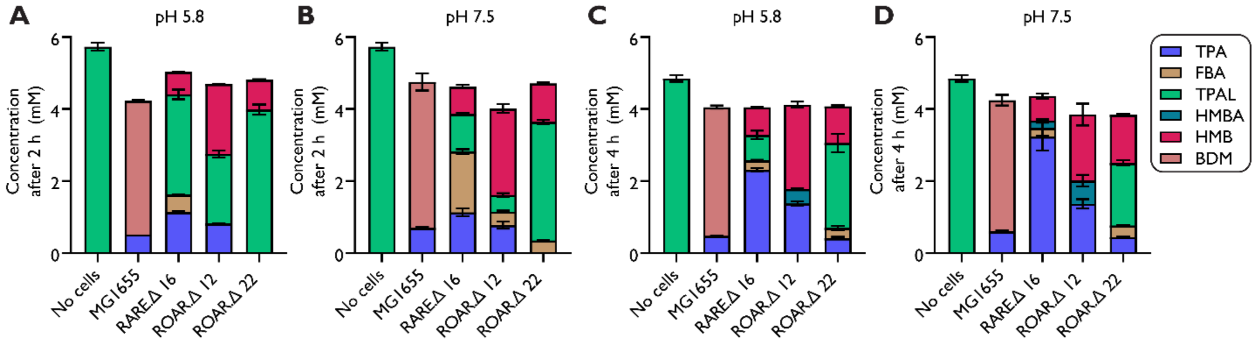


**Fig. S4. pH effect on TPAL stability resting cells.**

*E. coli* resting cells were cultured in LB media at 37 °C until mid-exponential phase, then dropped to 18 °C overnight for 18 h. The cells were washed and then resuspended in the 200 mM HEPES at (**A**, **C**) pH 5.8 and (**B**, **D**) pH 7.5 at 50 mg wet cell weight per mL. We then supplied 5 mM MHET-ald to MG1655, RARE.Δ16, ROAR.Δ12, and ROAR.Δ22 cells, and we measured the stability at (**A**, **B**) 2 h and (**C**, **D**) 4 h.

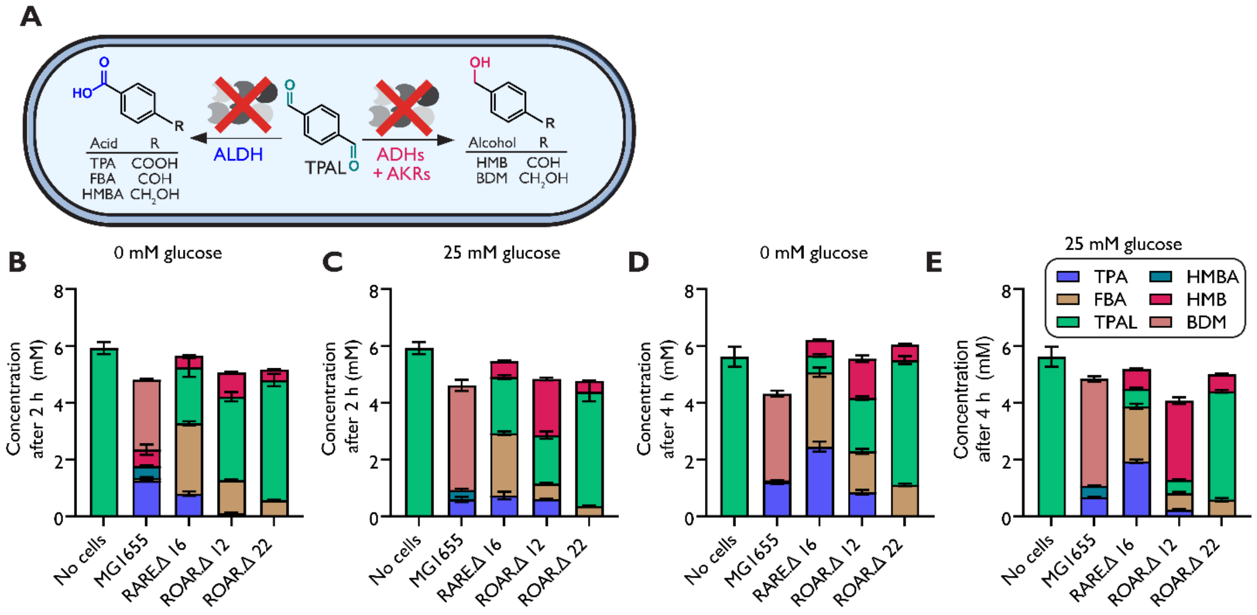


**Fig. S5. TPAL stability in resting cells.**

(**A**) *E. coli* resting cells were cultured in LB media at 37 °C until mid-exponential phase, then dropped to 18 °C overnight for 18 h. The cells were washed in 200 mM HEPES (**B**, **D**) without or (**C**, **E**) with 25 mM glucose at pH 7.5 and then resuspended in the same buffer at 50 mg wet cell weight per mL. We then supplied 5 mM MHET-ald to MG1655, RARE.Δ16, ROAR.Δ12, and ROAR.Δ22 cells, and we measured the stability at (**B**, **C**) 2 h and (**D**, **E**) 4 h.

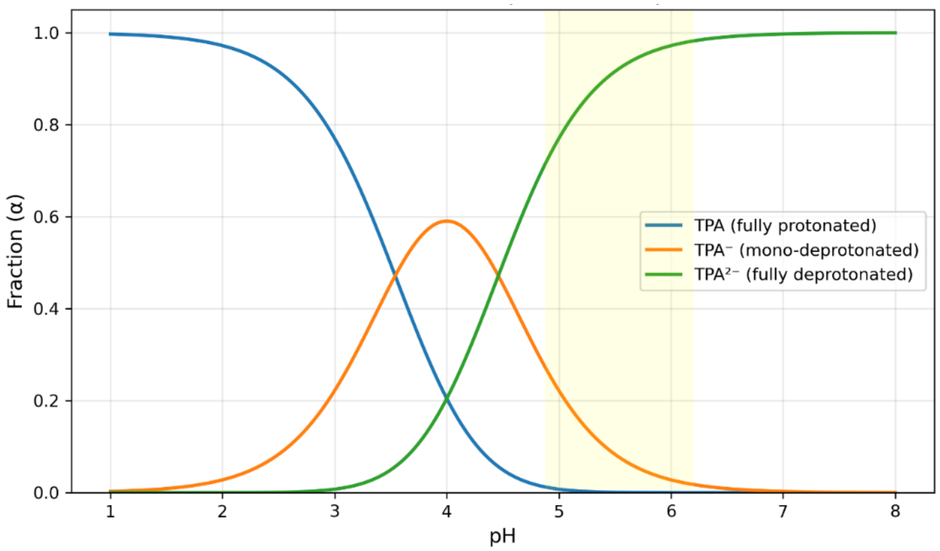


**Fig. S6. Predicted fraction of protonation states of TPA.**
The fraction of protonated species of TPA as related to pH was calculated using the Henderson–Hasselbalch relationship and the values of $pK_{a1} = 3.54$ and $pK_{a2} = 4.46$. The pH range where TPA import was observed and the trends of TPA reduction observed are consistent with the possibility that the mono-protonated species is actively imported by a native *E. coli* transporter. The fully protonated species precipitates in aqueous media.

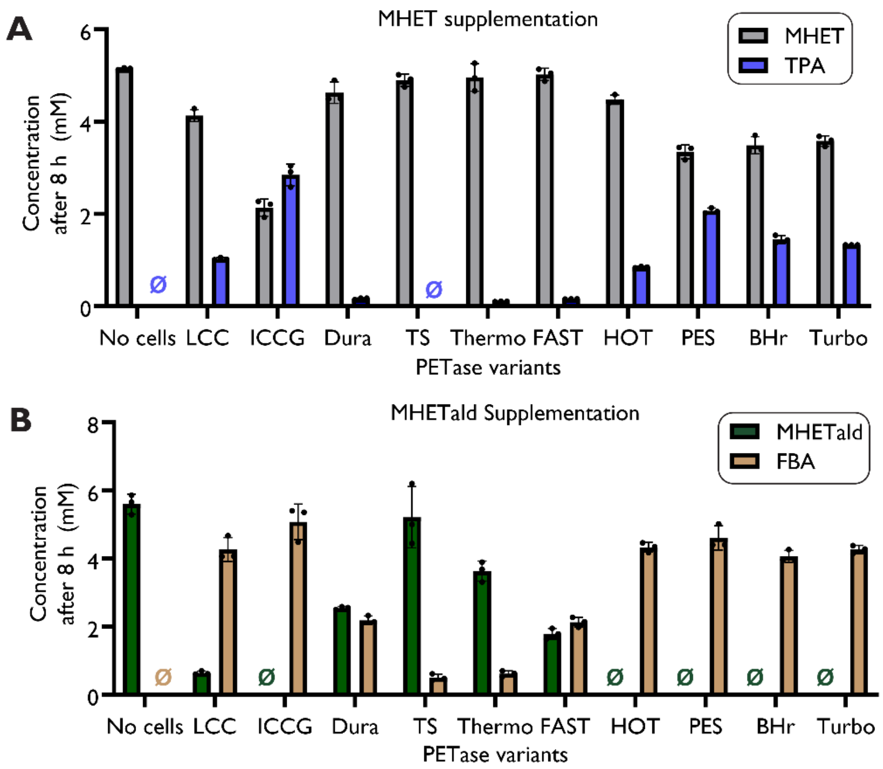


**Fig. S7. Evaluating the PETase specificity on PET deconstruction products**. Supplementation of 5 mM (**A**) MHET or (**B**) MHET-ald to RARE.Δ16 cell expressing PETase variants in MOPS media at pH of 7.4. Endpoint concentrations were measured after 8 h.

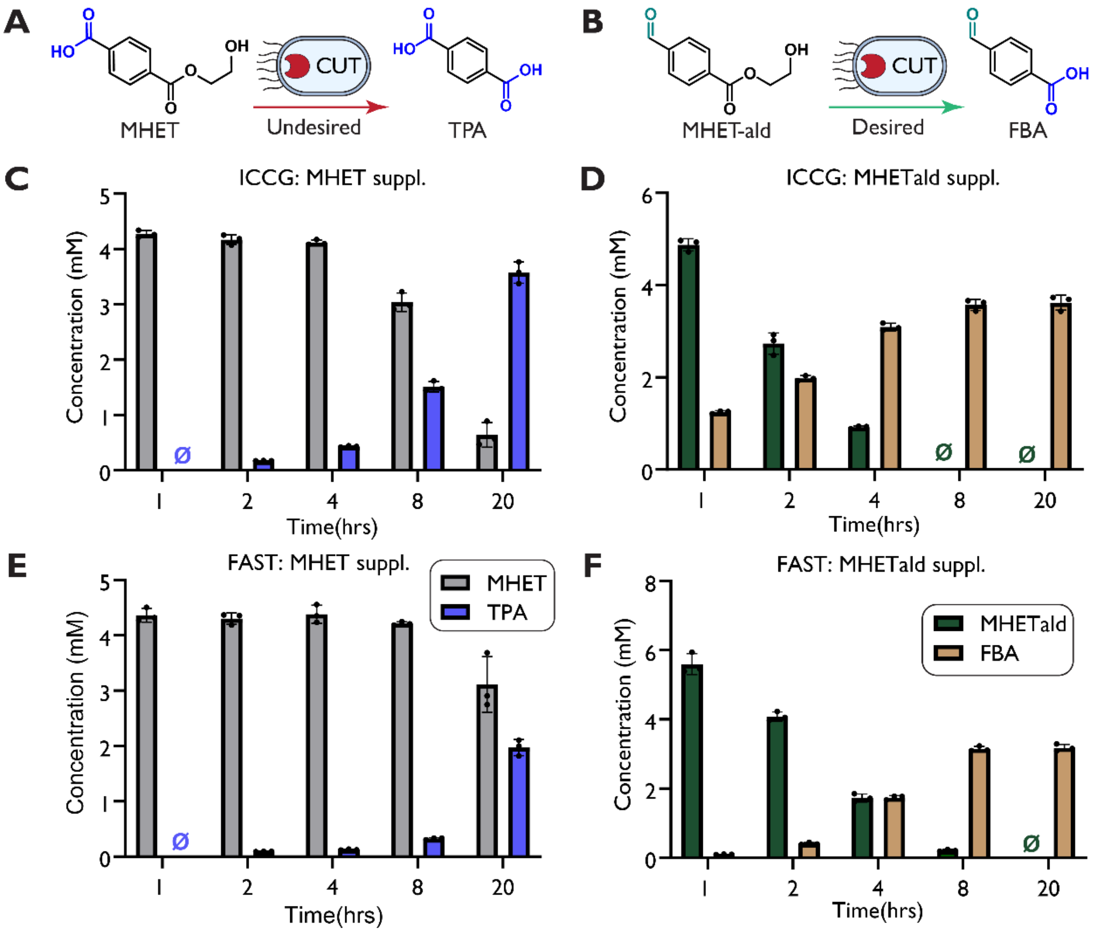


**Fig. S8. Time course of ETase activity on MHET and MHET-ald**

Evaluating the ETase activity of a TPA selective (ICCG) and a MHET selective (FAST) PETase on (**A**) MHET and (**B**) MHET-ald. Supplementation of (**C**, **E**) MHET or (**D**, **F**) MHET-ald to RARE.Δ16 cell expressing ICCG (**C**, **D**) or FAST (**E**, **F**) in MOPS media at pH of 7.4. Concentrations were measured for 20 h with timepoints taken at 1, 2, 4, 8, and 20 h.

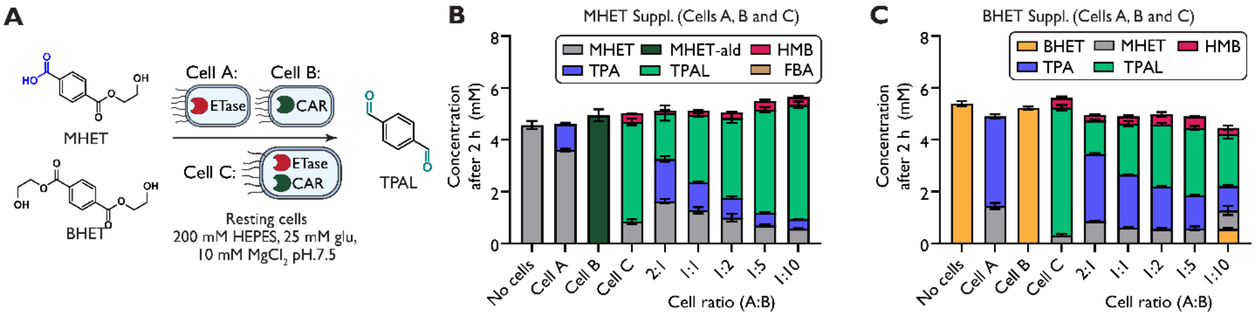


**Fig. S9. TPAL production in resting cells.**
(**A**) Depiction of resting cell TPAL production using FAST and MaCAR with MHET and BHET as a starting substrate. TPAL biosynthesis using resting cells of FAST (Cell A), MaCAR (Cell B) and FAST + MaCAR (Cell C) with (**B**) MHET and (**C**) BHET as starting substrates.

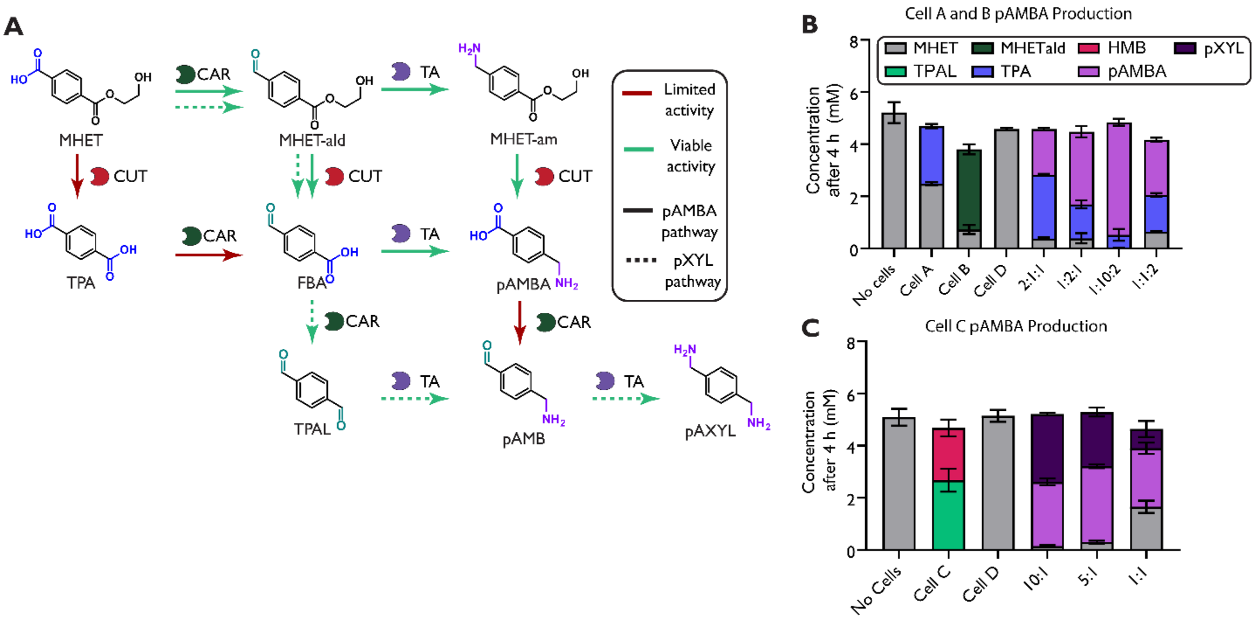


**Fig. S10. pAMBA production in resting cells.**
(**A**) Potential routes for amine production using FAST, MaCAR and CvTA from MHET as a starting substrate. (**B**) Resting cells of FAST (Cell A), MaCAR (Cell B) and CvTA (Cell D) were added to reaction mixtures with MHET at varying cellular ratios. (**C**) Resting cells of FAST and MaCAR (Cell C) and CvTA (Cell D) were added to reaction mixtures with MHET at varying cellular ratios.

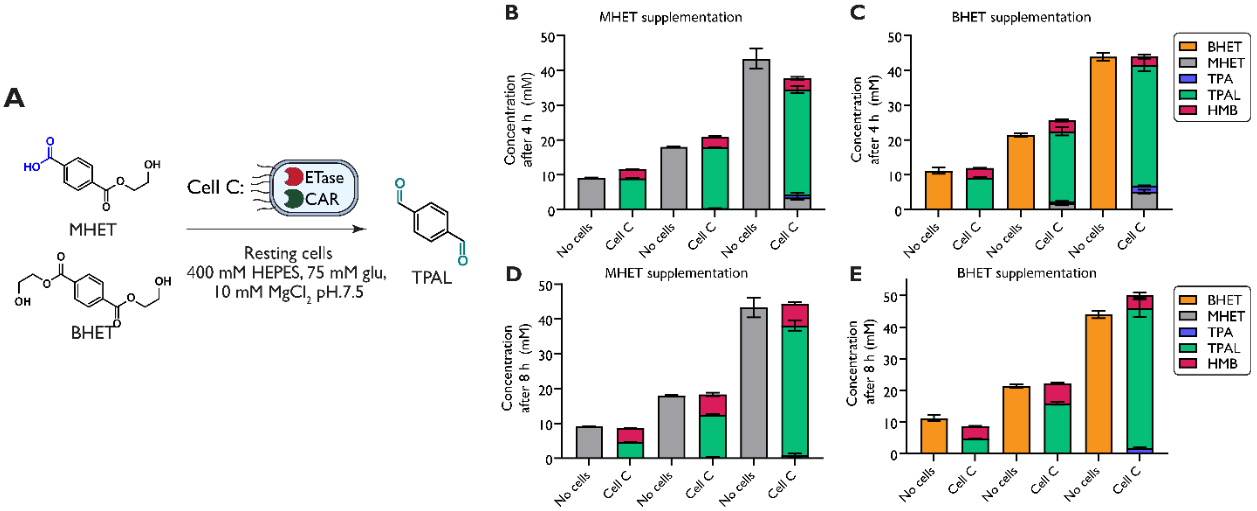


**Fig. S11. High substrate loading TPAL production.**

(**A**) Resting ROAR.Δ22 cells containing overexpressed FAST and MaCAR (Cell C) at higher substrate loading (10, 20 and 40 mM) of (**B**, **D**) MHET or (**C**, **E**) BHET were incubated at 30 °C with 400 mM HEPES, 10 mM MgCl2, and 75 mM glucose. Reactions were sampled at (**B**, **C**) 4 h and (**D**, **E**) 8 h.

### References


1. Selvam, E., Luo, Y., Ierapetritou, M., Lobo, R. F. & Vlachos, D. G. Microwave-assisted depolymerization of PET over heterogeneous catalysts. *Catal. Today* **418**, 114124 (2023).
2. Andini, E., Bhalode, P., Gantert, E., Sadula, S. & Vlachos, D. G. Chemical recycling of mixed textile waste. *Sci. Adv.* **10**, eado6827 (2024).
3. Gopal, M. R. *et al.* Reductive Enzyme Cascades for Valorization of Polyethylene Terephthalate Deconstruction Products. *ACS Catal.* 4778–4789 (2023) doi:10.1021/acscatal.2c06219.
4. Kunjapur, A. M., Tarasova, Y. & Prather, K. L. J. Synthesis and Accumulation of Aromatic Aldehydes in an Engineered Strain of Escherichia coli. *J. Am. Chem. Soc.* **136**, 11644–11654 (2014).
5. Butler, N. D., Anderson, S. R., Dickey, R. M., Nain, P. & Kunjapur, A. M. Combinatorial gene inactivation of aldehyde dehydrogenases mitigates aldehyde oxidation catalyzed by E. coli resting cells. *Metab. Eng.* (2023) doi:https://doi.org/10.1016/j.ymben.2023.04.014.
6. Dickey, R. M., Jones, M. A., Butler, N. D., Govil, I. & Kunjapur, A. M. Genome engineering allows selective conversions of terephthalaldehyde to multiple valorized products in bacterial cells. *AIChE J.* **69**, e18230 (2023).
7. Gopal, M. R. *et al.* Reductive Enzyme Cascades for Valorization of Polyethylene Terephthalate Deconstruction Products. *ACS Catal.* **13**, 4778–4789 (2023).
